# Alveolar macrophages initiate the spatially targeted neutrophil recruitment during nanoparticle inhalation

**DOI:** 10.1101/2024.11.13.623349

**Authors:** Qiongliang Liu, Lin Yang, Chenxi Li, Qiaoxia Zhou, Lianyong Han, Andreas Schröppel, David Kutschke, Judith Secklehner, Ali Önder Yldirim, Dagmar Zeuschner, Leo M. Carlin, Markus Sperandio, Otmar Schmid, Tobias Stoeger, Markus Rehberg

**Affiliations:** Institute of Lung Health and Immunity (LHI), Comprehensive Pneumology Center (CPC), Helmholtz Center Munich; Member of the German Center for Lung Research (DZL), Munich, Germany; Department of Thoracic Surgery, Shanghai General Hospital, Shanghai Jiao Tong University School of Medicine, Shanghai, 200080, China; Cancer Research UK Scotland Institute, Glasgow, UK; School of Cancer Sciences, University of Glasgow, Glasgow, UK; Electron Microscopy Facility, Max Planck Institute for Molecular Biomedicine, Muenster, Germany; Walter Brendel Centre of Experimental Medicine, Biomedical Center, Institute of Cardiovascular Physiology and Pathophysiology, Ludwig-Maximilians-Universität München, Planegg-Martinsried, Germany

## Abstract

Exposure to air pollution, including nanoparticles (NPs), is a major health concern associated with various diseases, triggered by subtle inflammatory responses in the lung. To investigate the dynamic immune response in vivo, lung intravital microscopy (L-IVM), was used to analyze the behavior of alveolar macrophages (AMs) and neutrophils, combined with ventilator-assisted inhalation of nebulized NPs in mice. Inhalation of fluorescent quantum dot NPs (cQDs) and soot-like carbon black NPs (CNPs, ambient pollutants), led to rapid spatially focused recruitment of neutrophils near alveolar deposited NPs. Neutrophil recruitment was initiated by NPs uptake by AMs, dependent on AM motility and AM NP surface recognition. Prior airway application of neutralizing antibodies against alveolar ICAM-1 and LFA-1, leading to reduced AM motility, inhibition of C5aR1 and FcγRI receptor mediated NPs uptake by AMs, as well as neutralizing of TNFα and application of a cellular degranulation inhibitor, abolished the early immune response induced by NPs. Overall, our data demonstrates the crucial role of AM activity (migration, phagocytosis, cytokine release) in the rapid and site-specific recruitment of neutrophils during the early phase of particle inhalation, suggesting these processes to be key events in mounting the immune response upon NP inhalation in the lung.

**Graphical abstract:** 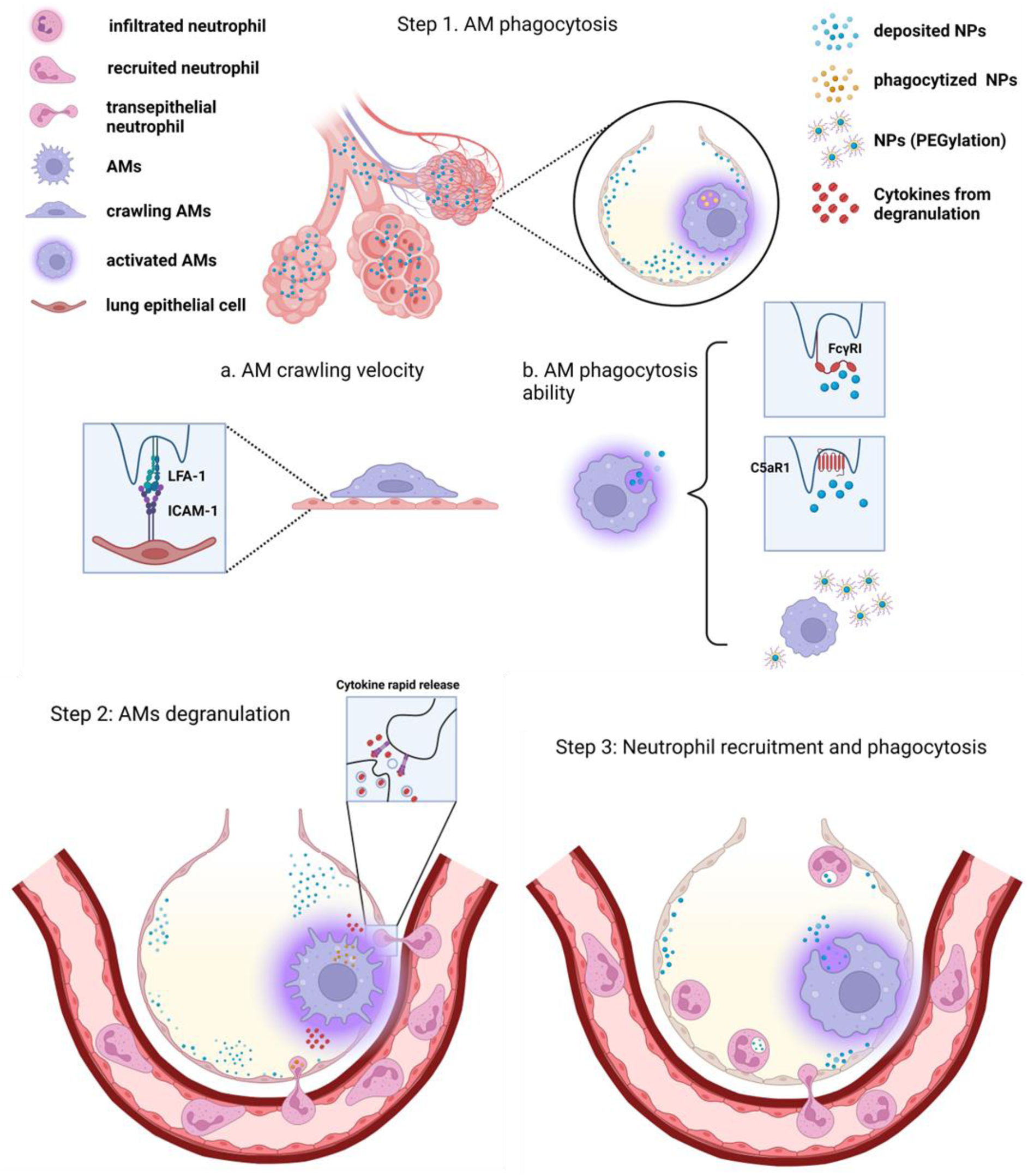

## Introduction

Respiratory and cardiovascular diseases are closely associated with air pollution, particularly due to airborne particulate matter, including fine particles (PM 2.5 with a diameter ≤ 2.5 µm) and ultrafine particles or nanoparticles (PM 0.1 with a diameter ≤ 0.1 µm), which contribute to increased mortality rates^1^. Owing to their nanoscale size, these particles are highly efficient at penetrating deep into the lungs, depositing in the fragile alveolar regions, and posing a greater health risk compared to larger particles due to their high surface area-to-mass ratio^2^. Inhalation of manufactured and incidental nanomaterials and nanoparticles (NPs: 1-100 nm) also largely contributes to acute respiratory distress syndrome and the development of chronic lung diseases^3,4^. Acute and transient pneumonitis is the most common reaction that can be triggered in vivo by almost any inhaled nanomaterial, depending on the dose delivered to the airway barrier. Although the most biologically relevant dose metric for the acute inflammatory response to inhaled poorly soluble low toxicity particles has been identified as the surface area of deposited NPs^5^, there is still a lack of understanding of the underlying molecular processes triggered by inhaled substances such as NPs, which is crucial for the development of effective preventive and therapeutic strategies.

Carbonaceous NPs represent an important proportion of ambient, urban particles, and pulmonary exposure to low doses has been shown to induce rapid proinflammatory responses in the lung including the release of pro-inflammatory cytokines^6–8^. Mechanistic information on the causal chain of the toxicological response cascade is furthermore needed for the development of the so-called Adverse Outcome Pathway (AOP) framework, a pragmatic approach facilitating non-animal testing strategies for the safety assessment of engineered NPs^9^. The AOP concept describes linkages between a molecular initiating event, to subsequent key events (KEs) leading to the adverse outcome, such as chronic lung disease or cardiovascular disease. Noteworthy several aspects of the local, inflammatory response to NPs have been described as KEs of various AOPs, and the particle-associated lung disease developed AOP is centered around the accumulation of neutrophils in the airspace^10,11^. It is well-known that tissue-resident macrophages and attracted polymorphonuclear neutrophils, as important parts of the innate immune system, act rapidly and non-specifically to protect the body upon any harmful lung exposures (e.g., pathogens or carbon-based particles)^12–15^. Although the immune response elicited to pathogens via specific receptors is well understood, a comprehensive understanding of the early events that trigger the innate immune response following NP inhalation, remains lacking, despite extensive research on the respiratory toxicity of NPs.

Previous studies on inhalation toxicology and immune responses following NP exposure have often relied on static time points for analysis, using cultured cells or animal models^16–18^. However, most experimental approaches lack the ability to capture both the in vivo biodistribution of NPs and the dynamic cellular cycles that precede and trigger specific biological effects, particularly in the early phases of NP exposure. This limitation might contribute to the fact that the sequence of events, in particular the contribution of tissue-resident macrophages in triggering spatially and temporally defined neutrophil recruitment, is still unclear.

Alveolar macrophages (AMs), situated in close contact with the lung epithelium in the alveolar space, play a crucial role as the first line of defense against inhaled pathogens^19^ and air-pollution-derived particles. AMs have been linked to the initiation of lung inflammation caused by air pollution particles^20^, however sensing of microbial components such as endotoxin by specific pattern recognition receptors such as TLRs, might modulate the inflammatory responses to ambient PM, and the role of AMs as initiating cell for sterile and pyrogen free particles might be different^7^. To resolve these contradictions, it is essential to investigate the alveolus during particle inhalation and study the ensuing cellular events in real-time.

The dense capillary network of the lung maximizes the direct contact between air and blood, enhancing oxygen exchange efficiency^21^. The enormous surface of the air-blood barrier, however, also increases susceptibility to inhaled NPs and pathogenic infections. Neutrophils, are the first responders recruited to sites of injury or infection, marking the onset of acute inflammation, but they also play a role in its resolution. In the lung, neutrophils amplify the immune response by infiltrating the alveoli, releasing chemokines, and activating downstream signaling pathways^21–23^.

Lung intravital microscopy (L-IVM) has been successfully employed to study neutrophil dynamics in microvessels in the alveolar region of murine lungs^24,25^ and has recently been extended to the analysis of AM functions, whereby AM crawling in and between alveoli has been demonstrated^26^. The autonomous defense behavior of AMs (intra alveolar patrolling and phagocytosis) seems to be closely linked to NP-induced neutrophilic inflammation. However, the spatio-temporal dynamics of AM activity and NP-induced neutrophil recruitment from the pulmonary microvasculature to the alveolar compartment during the early phase of NP-induced lung response remain elusive. Additionally, the molecular pathways underlying AM adhesion, migration, and NP recognition triggered by inhaled NPs are yet unknown.

To illuminate the real-time pulmonary innate immune response in the early stages of inhaled NP exposure in the peripheral alveolar region of the murine lung, we employ cutting-edge L- IVM combined with a ventilator-assisted aerosol delivery system. We used fluorescent quantum dot (QD) NPs to monitor particle deposition and uptake and carbon black (soot) NPs (CNP) as a surrogate for ambient particles. This approach allowed for the first time concomitant real-time observation of NP-aerosol deposition in the lung as well as the corresponding behavior of lung resident phagocytes and recruitment of leukocytes from the pulmonary microcirculation. Moreover, we investigated the molecular mechanisms governing the innate immune response by using various functional blocking antibodies / inhibitors and by altering the surface properties of inhaled NPs.

The presented data indicate a close relation between AM activity (phagocytosis, migration) and the rapid and NP site-specific recruitment of neutrophils during the early phase of NP inhalation (< 60 min), demonstrating a specific role of AMs in mounting the immune response upon NP inhalation.

## Results

To observe in real-time the deposition of inhaled particles in the lungs of living mice and simultaneously study the immune response in the alveolar region of the lungs, we combined a ventilator-assisted aerosol inhalation system with lung intravital microscopy (L-IVM)^27^ (Figure 1a, b). To characterize NP lung deposition dynamics, we utilized fluorescent carboxyl quantum dot NPs (cQDs, diameter: 15 nm), which have been extensively used to study adverse in vivo effects and cellular uptake of NPs^28–31^. The nebulizer used for ventilator-assisted aerosol delivery generates aerosol droplets (2.5-4.0 μm) containing NP suspensions. The NP doses (cQDs and in the following CNPs) were chosen such that they induce a comparable level of neutrophil recruitment in the lung.

**Figure 1.**
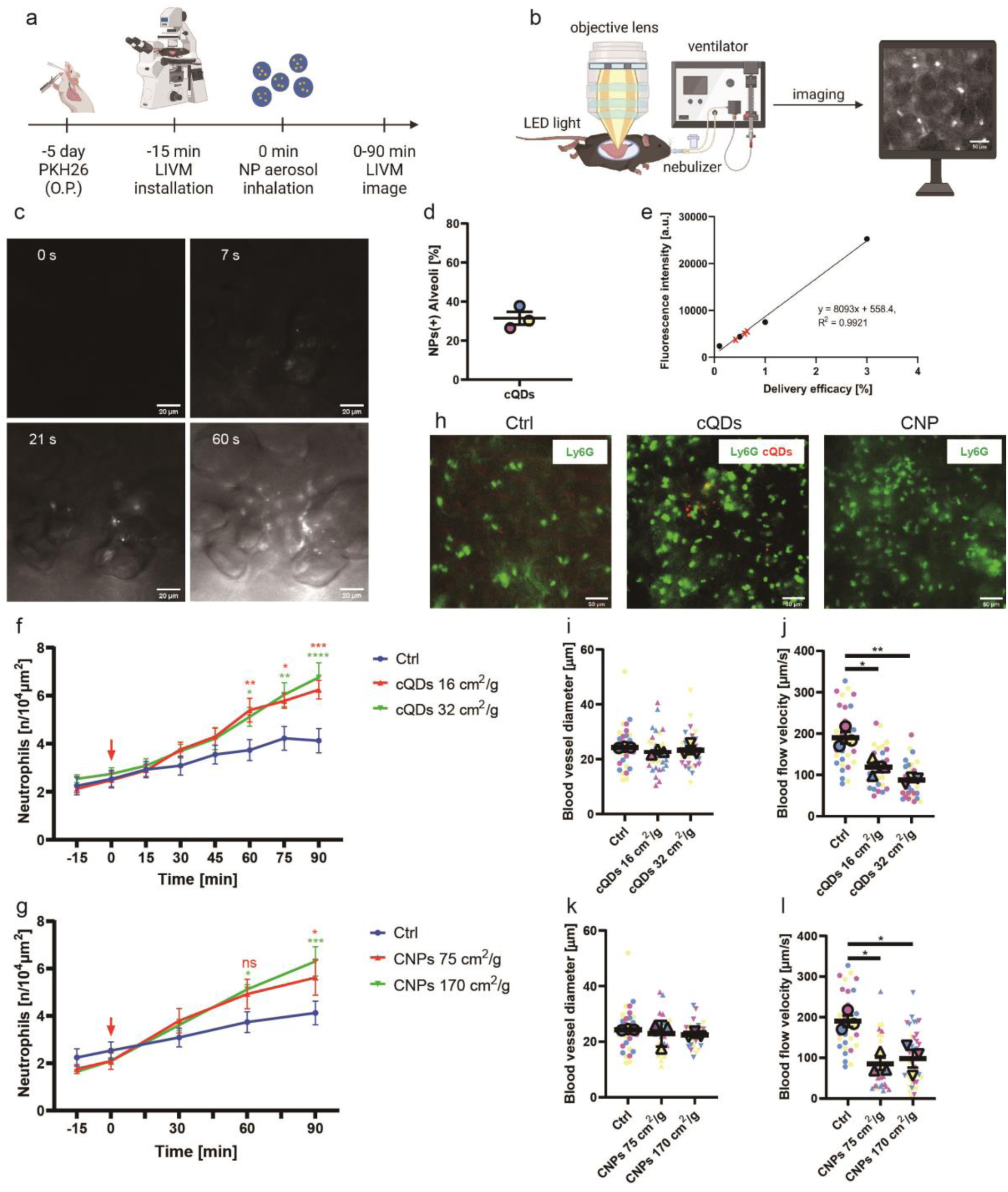
NP inhalation causes rapid recruitment of neutrophils in the pulmonary microcirculation, with associated hemodynamic changes. (a) Schematic representation of NP inhalation experiments, including pretreatment for alveolar macrophage imaging. (b) Schematic representation of lung intravital microscope (L-IVM) in mice connected to a mechanical ventilator equipped with a nebulizer for NP aerosol inhalation. (c) Inhaled cQDs were rapidly detected by L-IVM as distinct fluorescent spots, reflecting cQDs containing aerosol droplets (yellow arrows) in single alveoli. Over inhalation time the number of deposited cQDs spots and fluorescence intensity increased. Scale bars: 50μm. (d) 5 min after cQDs inhalation, quantification of cQD positive alveoli by L-IVM (16 cm^2^/g (geom. surface area of NPs / mass-lung)), 7 field of views (FOVs) / mouse; each FOV contains 20-30 alveoli, n=3 mice. The mean values of the individual mice are shown as larger circles of the same color. (e) cQDs delivery efficacy to the lung was calculated according to the linear fluorescence intensity-standard curve (R^2^ > 0.99) obtained from blank lung homogenates spiked with known cQDs doses normalized to the aerosolized cQD dose (black circles). Red marks indicate the QDs fluorescence intensities detected in mouse lung homogenates of 3 independent inhalation applications. (f) Quantitative analysis of neutrophil numbers after inhalation of two different doses of cQDs. NPs inhalation starting at 0 min, indicated by red arrow. Inhalation typically lasted 1 min for QDs. n=6 mice/group. (g) Quantitative analysis of neutrophil numbers after inhalation of two different doses of CNPs. NPs inhalation starting at 0 min, indicated by red arrow. Inhalation typically lasted 5 min for CNPs. n= 4-6 mice/group. (h) Representative L-IVM images of neutrophils (anti-Ly6G mAbs, i.v., green) at 90 min after inhalation of vehicle (control), cQDs (16 cm^2^/g; red) and CNPs (75 cm^2^/g). Scale bars: 50 μm. (i) and (k) Diameter of alveolar microvessels in the observation area are not affected by inhalation of cQDs and CNPs, respectively. Each data point represents the diameter of one microvessel. The means of the respective mouse are displayed as larger circles and triangles of the same color (n=3). (j) and (l) Blood flow velocities in the pulmonary microcirculation of QD- and CNT-exposed mice are reduced, respectively. Determination of blood flow velocity by tracking trajectories of i.v. injected fluorescent beads (at t=90 min, bead diameter = 1 µm). Each data point represents the velocity of one fluorescent bead/vessel, 10 beads tracked per mouse (n=3). Data are presented as Means ± SEM, f,g,i,,j,k,l: One-way ANOVA test.

NPs reached peripheral alveoli seconds after the onset of inhalation and were immediately detected as distinct fluorescent spots by L-IVM (Figure 1c). Notably, not all alveoli in the field of views exhibited deposited cQDs. At the administered dose of 16 cm^2^/g (geometric NP surface area / mass-lung) used in this study, 31.47±2.2 % of alveoli in the recorded field of views received cQDs (Figure 1d). To quantify the lung deposited cQD-NP dose after ventilator- assisted nebulization of 20 µl of a 4 µM cQD suspension, mice were sacrificed immediately after NP application and the QD fluorescence in the lung tissue was quantified via spectrofluorometry in lung homogenate. The cQDs deposition efficacy was determined as 0.53±0.10 % (Mean±SEM; Figure 1e).

To investigate the potential inflammatory effects of cQD-NPs in the alveolar region, L-IVM was employed to observe and quantify neutrophil recruitment upon ventilator-assisted cQDs inhalation in pulmonary microvessels. For this, neutrophils were immunolabeled in the vascular compartment by intravenously (i.v.) injected fluorescent anti-Ly6-G antibodies 10 min prior to imaging (Figure 1b). Already at 30 min upon inhalation of a lung deposited dose of 16 cm^2^/g cQDs, an increased number of neutrophils, as compared to the control group, was determined in the observation areas, which became significant at 60 min. Interestingly, increasing the deposited dose (from 16 cm^2^/g to 32 cm^2^/g) of cQDs did not cause a significant difference in the numbers of recruited neutrophils, as determined by semi-automated quantitative detection of fluorescent neutrophils (Figure 1f). To determine whether leukocyte recruitment dynamics obtained for cQDs are comparable to the inflammatory response induced by inhalation of a bioequivalent dose of carbon nanoparticles (CNP), a common component of urban air pollution^7^, mice were exposed to calculated deposited CNP doses of 75 cm^2^/g and 170 cm^2^/g by ventilator-assisted inhalation. Indeed, 60 min after CNP inhalation, a large number of neutrophils accumulated in the lungs of mice receiving 75 cm^2^/g or 170 cm^2^/g CNP via inhalation (Figure 1g, h), with cell counts comparable to those observed in cQDs-exposed animals (Figure 1f, h). From Fig 1f and 1g, it is apparent that the onset of neutrophil recruitment is independent of dose, while the rate of recruitment may be dose dependent at later time points which is seen as a non-significant trend in the respective 90 min data.

Inflammation can potentially cause changes in pulmonary microvascular hemodynamics that affect blood flow velocity and shear rates^25^. To assess the impact of NP inhalation on the alveolar microcirculation, blood flow velocities in the lungs of cQDs-exposed mice were measured by tracing of i.v. injected fluorescent microbeads. Blood flow velocities were decreased at 90 min after inhalation of both cQDs and CNPs (Figure 1j, l). Pulmonary microvessel diameters were not altered among the experimental groups (Figure 1i, k), excluding local vasoconstriction or vasodilation as a cause for NP inhalation induced blood flow velocity alterations.

Taken together inhalation of cQDs as well as CNPs caused rapid neutrophil recruitment and blood flow velocity reduction in the pulmonary microcirculation.

Differential cell analysis in bronchoalveolar lavage (BAL) fluid was applied to further define inflammatory responses caused by cQDs or CNP inhalation (16 cm^2^/g cQDs, 170 cm^2^/g CNP). A low number of neutrophils was detected 2 h after inhalation of cQDs, whereas neutrophils were almost undetectable in control animals (Figure S1a). Likewise, in the CNP-exposed group, neutrophil numbers were elevated, although not significant (Figure S1b). Compared with the control group, the amounts of total cells, AMs and lymphocytes were similar in the cQDs- as well as the CNP-exposed group (Figure S1a, b). At 24 h upon cQDs and CNP-inhalation, a substantial influx of neutrophils (5-10% of total BAL cells) into the airspace occurred, contrasting with almost undetectable levels in BAL of control mice (Figure S1c, d).

Taken together, BAL analysis confirmed an acute inflammatory response induced by cQDs and CNPs, marked by a notable accumulation of neutrophils at 2 h and 24 h after exposure.

### Neutrophils are recruited in pulmonary microvessels in close proximity to the sites of cQD-NP deposition

Close inspection of long-term L-IVM recordings of neutrophil dynamics upon cQD inhalation suggested that neutrophils preferentially arrested in microvessels near the alveolar deposited cQDs, where they often exhibited a probing/crawling behavior (Figure 2a). Therefore, the number of neutrophils close to deposited cQDs was quantified in square areas (100 μm side length) centered around cQDs ‘hotspot’ (Figure 2b) and compared to neutrophil counts in cQDs-deficient areas. Indeed, the number of neutrophils increased significantly and more rapidly in close proximity to cQDs accumulations, than in cQDs-deficient areas (Figure 2c).

**Figure 2.**
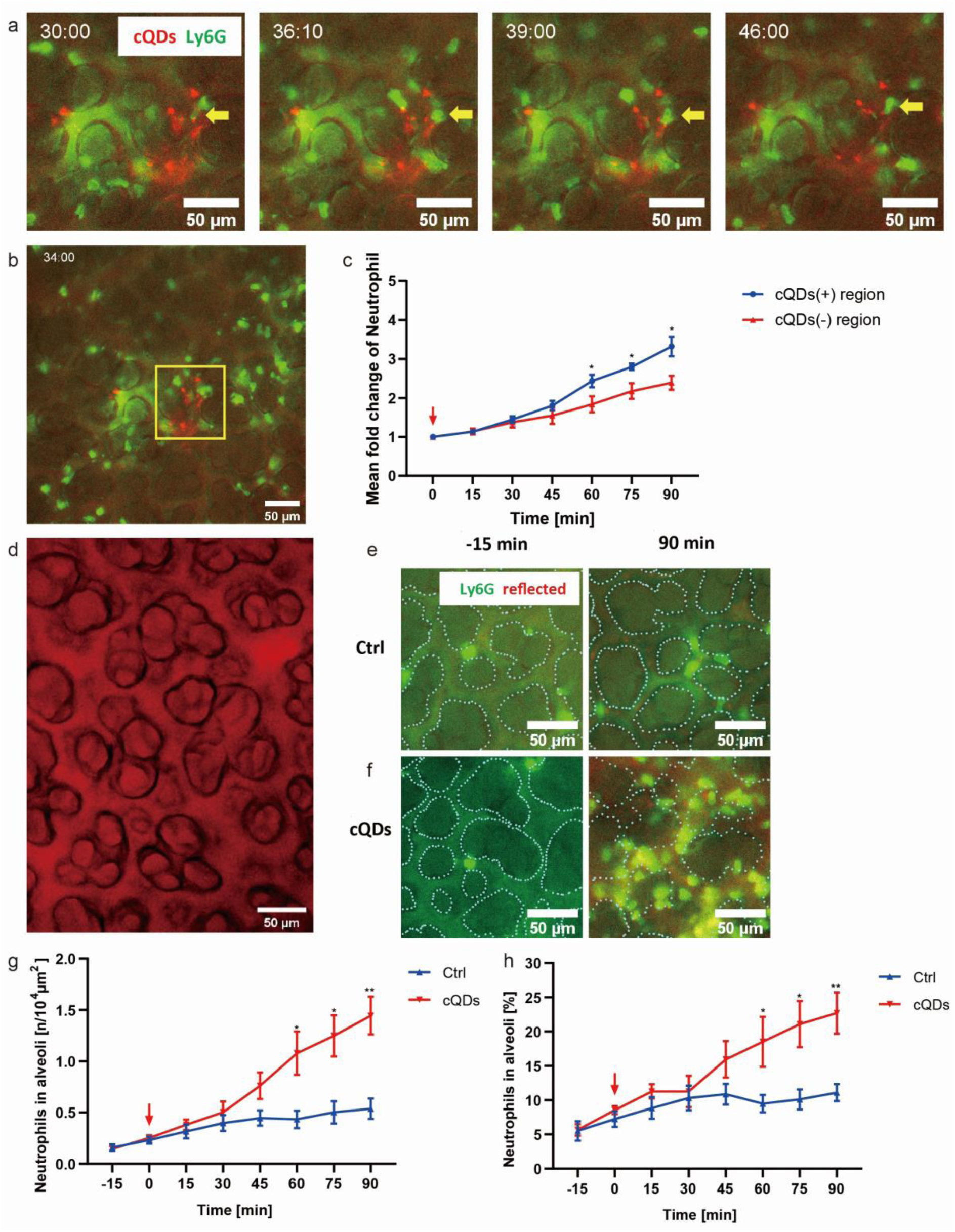
Neutrophil recruitment and infiltration into the alveoli in response to inhaled nanoparticles occur in close proximity to the site of NP deposition. (a) Time lapse of a cQD probing neutrophil (yellow arrow) in a cQD-abundant area. cQDs (Exposure dose: 16 cm^2^/g, red) exposed mice i.v. injected with anti-Ly6G to label neutrophils (green), Scale bar 50 μm. (b) A large number of neutrophils accumulate in alveolar regions with close proximity to deposited cQDs (yellow square 100 μm side length = 0.01 mm^2^). Scale bar 50 μm. (c) Neutrophil numbers over the time course of L-IVM were compared between cQDs abundant and cQDs deficient regions. Neutrophils in cQD abundant regions were analyzed in 0.01 mm^2^ squares centered on cQD hotspots as indicated in (b). Data are presented as mean fold change from the baseline as Means ± SEM, n=3 mice/group (7 FOVs per mouse), *: p < 0.05 by Two-way ANOVA test. (d) Visualization of the microstructure of the lung using L-IVM with reflected light. Alveoli and interstitial tissue, mainly containing microvessels, are clearly visible by reflected light (illumination/detection wavelength: 655 nm, Scale bars: 50 μm). The roundish areas represent the alveoli, which are surrounded by the microvascular bed. These structures are emphasized by the superimposed lines in (e) and (f). (e)-(f) L-IVM of control (e) and cQDs (16 cm^2^/g) exposed (f) mice injected with anti-Ly6G mAbs to label neutrophils (green) and alveolar structure was visualized by reflected light (red; 655 nm). Dotted line images represent alveolar boundaries (panel e: Ctrl group -15 min, panel g: Ctrl group 90 min, panel f: cQDs group - 15 min, panel h: cQDs group 90 min, Scale bars: 20 μm). (g) Neutrophil numbers infiltrated into alveoli over the time course of L-IVM after cQDs (16 cm^2^/g) inhalation as compared to controls. (h) Percentage of alveolar localized neutrophils of the total number of neutrophils after cQDs (16 cm^2^/g) inhalation and under control conditions. Data are presented as Means ± SEM in (c), (g) and (h), n=3 mice/group, *: p < 0.05 and **: p < 0.01 by Two-way ANOVA test. QDs inhalation starts at 0 min (red arrow).

During L-IVM, the lung structure can be clearly imaged using reflected light microscopy, thus the outlines of the alveoli are easily identifiable (Figure 2d, e, f). These imaging properties enable the discrimination of airspace and microvessels, and thus also the differentiation between alveolar and microvessel-localized neutrophils (Figure 2e, f). Already 30 min after inhalation of cQDs, an increased number of alveolar localized neutrophils could be detected compared to control mice, and the increase became significant at 60 min. (Figure 2g), with the proportion of alveolar localized neutrophils to total neutrophils increasing significantly faster than in the control animals over the course of time (Figure 2h). Overall, these data suggest that the influx of neutrophils into the airspace starts within 30-45 min after inhalation of cQDs.

Immunofluorescence staining of lung tissue sections of mice 90 min after cQDs inhalation, frequently identified lung neutrophils with internalized cQDs (Figure S2a), and in L-IVM images obtained 1 hour after cQDs inhalation, alveolar localized neutrophils were associated with cQDs, both indicating QDs uptake (Figure S2e). In agreement to that, analysis of bronchoalveolar lavage obtained 24 h after cQD inhalation showed that cQDs were not only phagocytosed by macrophages (23.7±1.2% of all BAL macrophages contain cQDs), but also by neutrophils (15.6±2.9% of all BAL neutrophils contain cQDs) (Figure S2b, c, d). Thus, these data indicate a contribution of neutrophils in the alveolar clearance of NPs.

### AMs exhibit enhanced patrolling and accumulate in NP-rich areas upon NP inhalation

AMs serve as main phagocytic cells in the alveolar lumen cavity, exhibiting robust capabilities in clearing inhaled pathogens and pollution-related particles (bacteria, viruses, NPs), as well as cellular debris and lung surfactant thereby maintaining lung homeostasis^32,33^. To facilitate the investigation of AM motility and NP uptake and relate these to neutrophil responses, AMs have been labeled by oropharyngeal aspiration of PKH-26 dye^26^. Our L-IVM images confirmed recent data from Neupane et al.^26^ that not every alveolus contains an AM (Figure 3a), with the ratio of AMs to alveoli being one to three (Figure 3b). This again emphasizes the need for AMs to patrol within and between alveoli to maintain homeostasis of the alveolar microenvironment^26^.

**Figure 3.**
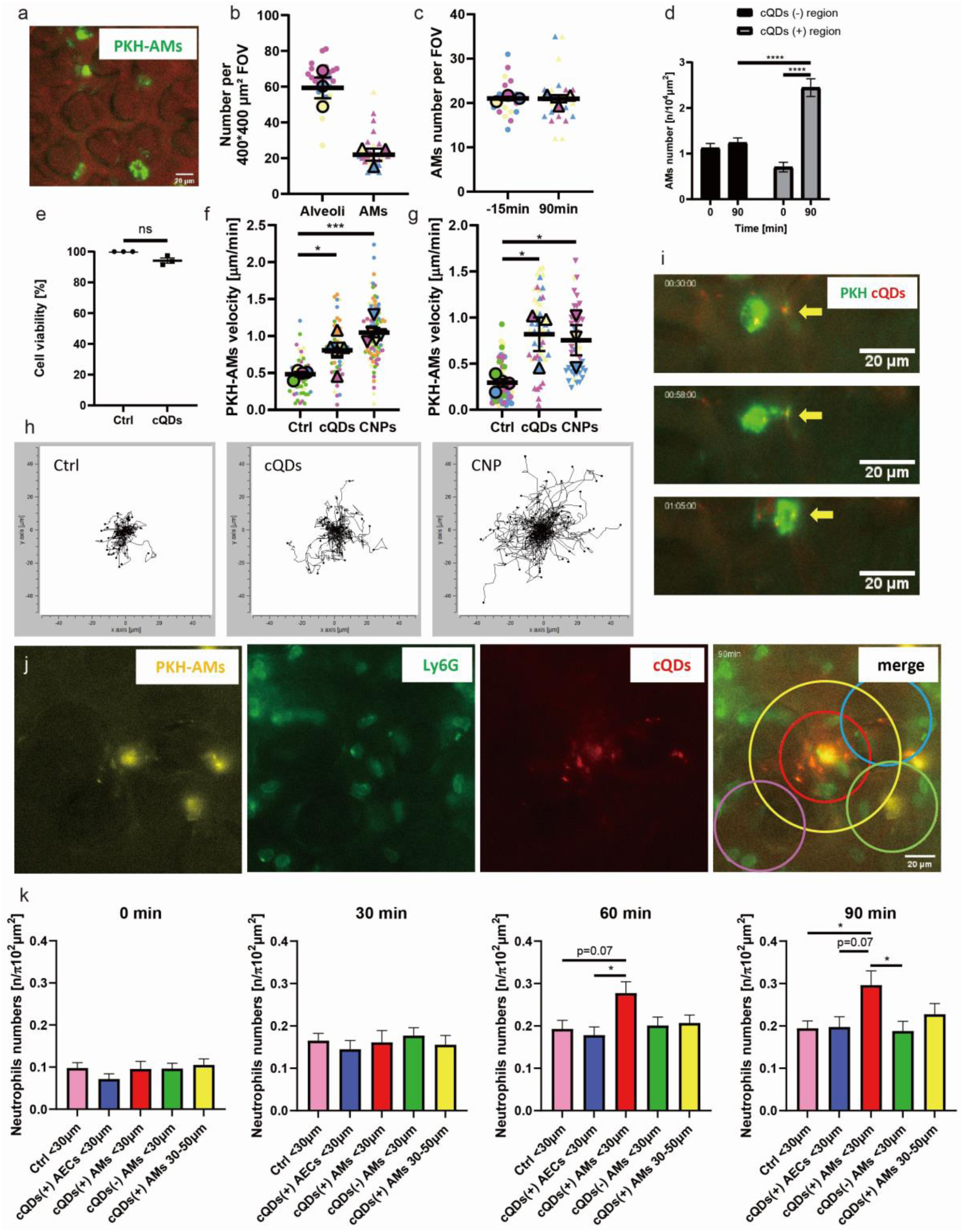
Alveolar deposition of NPs altered the patrolling speed of AMs and induced the recruitment of neutrophils that focally appeared around NP-containing AMs. (a) Representative L-IVM image of PKH-AMs (green), merged with reflected light image depicting lung microstructure (red). Scale bar: 20 μm. (b) Quantification of the ratio of AMs to alveoli in L-IVM images, 7-14 random region/mice, n= 3 mice/group. The mean values of the individual mice are shown as larger circles or triangles of the same color. (c) L-IVM quantification of PKH-AM numbers 15 min prior and 90 min after cQD exposure, 7 random regions/mice, n=3 mice/group. The mean values of the individual mice are shown as larger circles or triangles of the same color. (d) Upon 90 min after inhalation, more AMs are detected by L-IVM in cQDs-enriched areas (Diameter: 100μm), n=3 mice/group. (e) Cell viability of M-HS cells determined by WST assay after exposure to cQDs (8nM, this largely corresponds to the in vivo delivered dose) for 1 h, n=3. (f) and (g) average track velocity (µm/min) of AMs in different NP-exposed mice (f: 1 h) and (g: 24 h) during a 1 h L-IVM imaging session. Each symbol represents an individual randomly selected AM combined from 4-6 separate experiments, n=4- 6 mice/group. The mean values of the individual mice are shown as larger circles or triangles of the same color. (h) Spider plot of randomly chosen PKH-AM migration paths in the alveoli of mice after cQD and CNP exposures during a 1h L-IVM imaging session (0-1 h), 10-20 AMs / mice, 4-5 mice / group. (i) Time-lapse L-IVM of PKH-AM (green) crawling towards and engulfing cQDs (yellow, arrows). (j) L-IVM images after cQDs-inhalation (60min). Neutrophils (green) aggregated close to PKH-AMs (yellow) with internalized cQDs (red). The radii of the small and large circles are 30 μm and 50 μm, respectively. Center of circles: cQDs(+) PKH-AMs (red and yellow); cQDs(+) PKH-AMs (blue); cQDs(+) AECs (green) and control areas (pink). Scale bars: 20 μm. (k) Quantification of neutrophil accumulation in the areas indicated in (j), n=4. Data are presented as Means ± SEM. ns: p ≥ 0.05, *: p < 0.05, **: p < 0.01, ***: p < 0.001 and ****: p < 0.0001.

cQD inhalation did not affect PKH-AM cell numbers in alveoli in the analyzed field of views 15 min prior and 90 min post-cQD inhalation, as determined by L-IVM (Figure 3c). However, AMs accumulated over time close to deposited cQD hotspots (Figure 3d). Tracking of PKH- labeled AMs 60 min post-inhalation revealed that compared to control values, inhalation of cQDs, as well as CNPs, increased the migration velocity of AMs in the alveoli of the respective mice (Figure 3f, h). Importantly, even 24 h after cQDs and CNP exposure, AMs remained particularly active in the alveoli, exhibiting increased velocities compared to controls (Figure 3g). Increased AM patrolling might thus facilitate effective NP clearance, as observed within the first hour post-cQD inhalation, where AMs crawled towards and phagocytosed alveolar deposited cQDs (Figure 3i). We do not anticipate cytotoxic effects of cQDs, since incubation of a murine AM cell line, MH-S^34^, with cQDs did not affect cell viability (Figure 3e).

### Neutrophil recruitment is initiated by cQDs^(+)^ AMs

Having observed that neutrophil recruitment occurs in the vicinity of NP “deposition hotspots” (Figure 2c)^35–37^, we next used spatial segmentation to analyze whether AMs or alveolar epithelial cells contribute to initiating the local immune response at the alveolar level (mouse alveoli is approximately 30µm^38^ (Figure 3j).

Therefore, neutrophil numbers in circular regions of interest (r < 30 µm) centered around (i) PKH26-labeled AMs colocalized with cQDs (cQDs^(+)^ AM), (ii) AMs without cQDs (cQDs^(-)^ AMs), (iii) cQDs localized at/in alveolar walls / alveolar epithelial cells (AECs), or (iv) control regions (cQD as well as AM free) have been determined upon inhalation over time (Figure 3k).

From 60 min after cQD inhalation, significantly increased neutrophil aggregation appeared only around cQDs^(+)^ AMs compared to cell numbers detected adjacent to cQDs^(-)^ AMs and cQDs clusters localized at/in alveolar epithelial cells as well as control regions (Figure 3k). With increasing distance (from r < 30 µm to r = 30-50 µm), the tendency of neutrophils to gather in close proximity to the cQDs(+) AMs decreased (Figure 3k). Taken together, the data suggest that the cQD-induced spatially restricted recruitment of neutrophils is initiated by particle-laden AMs rather than the adjacent alveolar epithelial cells.

### Inhibiting cellular degranulation diminishes NP induced neutrophil recruitment

Considering that neutrophils accumulate close to NP-laden AMs, and macrophages are known to release pro-inflammatory mediators upon stimulation of pattern recognition receptors and receptor-mediated phagocytosis, we set out to investigate whether inhibited cellular degranulation (by application of cromolyn) is similarly involved in pulmonary neutrophil recruitment as previously shown post intravenous NP injection in skeletal muscle tissue^28^. Noteworthy, cromolyn has been suggested to inhibit AM stimulation via plasma membrane stabilization^39^.

Mice were pretreated by i.v. injection with 0.2 mg/kg (BW) cromolyn 30 min before NP inhalation (Figure 4a). Cromolyn-pretreatment prevented cQD- and CNP-induced neutrophil recruitment and neutrophil levels after NP inhalation matched those of the respective vehicle control group (Figure 4b, c). The anti-inflammatory effect of cromolyn was long-lasting, pretreatment with cromolyn prevented infiltration of neutrophils into the airspace for 24 h post- NP exposure, as determined by BAL analysis (Figure 4d, e). Accordingly, cromolyn pretreatment restored blood flow velocity in cQDs-exposed mice to control levels with no discernible impact on pulmonary microvessel diameters (Figure 4f).

**Figure 4.**
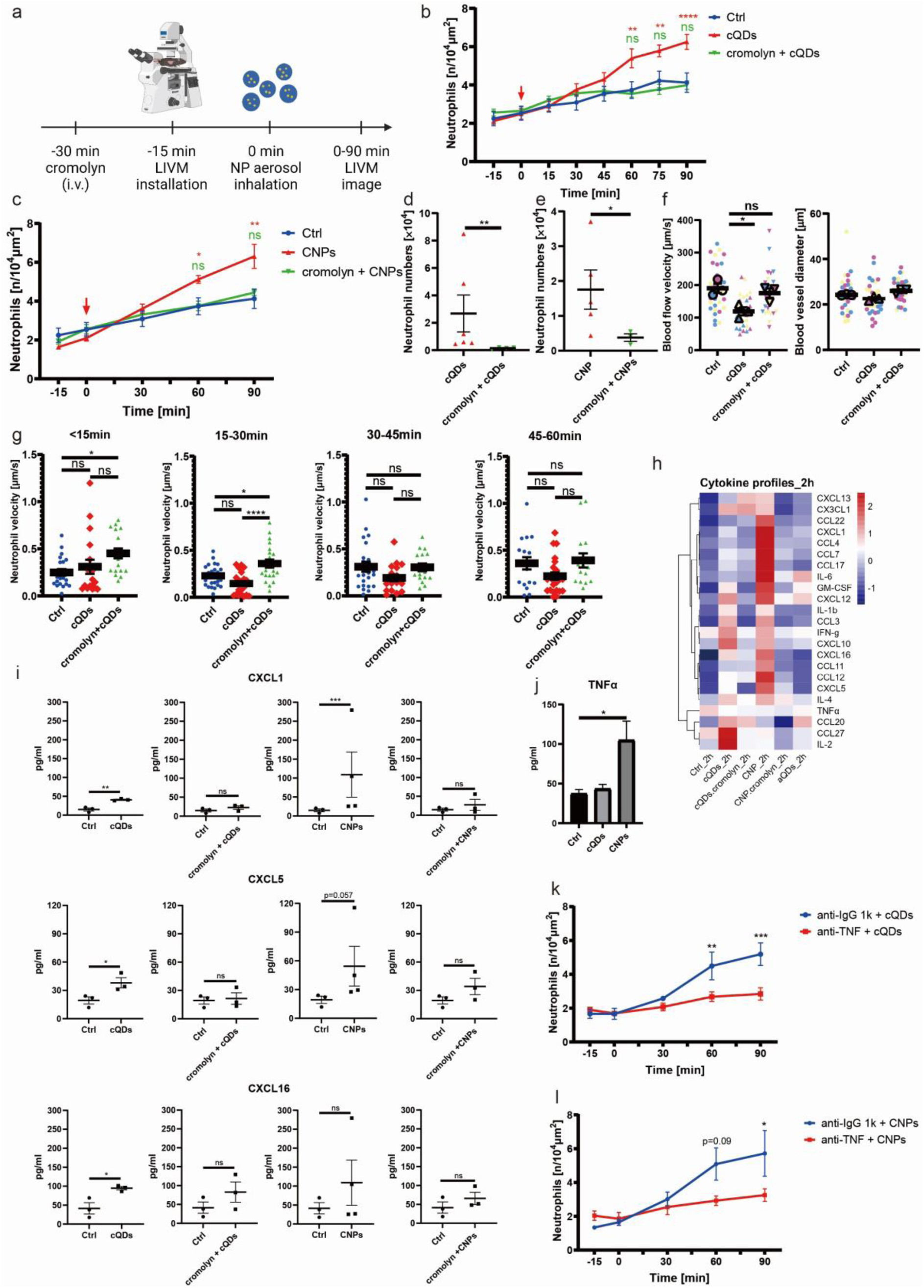
NPs-induced neutrophil recruitment is initiated by cellular degranulation. (a) Schematic of inhibiting cellular degranulation by cromolyn administration. (b)-(c) Quantification of neutrophil counts during L-IVM in mice receiving cromolyn (i.v. injection of 0.2 mg/kg BW) 30 min prior to NP inhalation of (b) cQDs, (c) CNP or vehicle, n=4-6 mice/group. (d)-(e) Neutrophil numbers detected in BAL cytospin samples of mice (pretreated with cromolyn or vehicle) at 24 h after cQDs (d) or CNP (e) inhalation. Panel d: cQDs, n=4 mice/ group and panel e: CNP, n=3 mice/ group. (f) 90 min after cQDs inhalation, blood flow velocity in the pulmonary microvessels of mice, which are treated with or without cromolyn 30 min prior to inhalation (left). Determination of blood flow velocity by tracking trajectories of i.v. injected fluorescent beads. Each data point represents the velocity of one fluorescent bead/vessel, 10 beads tracked per mouse, respective mean values are superimposed as larger circles and triangles of the same color (n=3). Blood vessel diameter in the fields of view of the different treatment groups (right). n=3 mice/group. (g) Crawling velocity of neutrophils with long residence time (>5 min) in the vicinity of cQDs-abundant areas (distance < 100 μm) in mice with or without cromolyn-pretreatment was analyzed in L-IVM recordings after inhalation of cQDs or vehicle-control. 6-7 neutrophils/mouse, 3 mice/group. (h) Heatmap representation of chemokine concentrations in BAL samples obtained 90 min after inhalation exposure of cQDs, aQDs and CNP in naive mice or pretreated with or without cromolyn, n=3-4 mice/group. (i) Individual alterations of cytokines. Cytokine concentrations were detected in mouse serum samples upon NPs inhalation for 2 h as well as under control conditions. (j) Concentrations of TNFα quantified by ELISA in BAL fluid from mice 2 h after CNP, cQDs and vehicle inhalation, n=3. (k)-(l) Alterations in the amounts of neutrophils over time under L-IVM were compared between mice pretreated with anti- TNFα mAb or isotype mAbs followed by cQDs (k) or CNP (l) inhalation, n=3 mice/group. NPs inhalation starting at 0 min, red arrow. Data are presented as Means ± SEM, ns: p ≥ 0.05, *: p < 0.05, **: p < 0.01 and ***: p < 0.001 by Two-way ANOVA test.

To further study the effect of cromolyn on cQDs-induced neutrophil recruitment, we analyzed the neutrophil crawling velocity along the microvessel walls. Following cQDs exposure, the speed of neutrophil movement along blood vessel walls near deposited cQDs (from 15 min) was decreased compared with the situation in control mice, again demonstrating the rapidity of the neutrophil response. Pretreatment with cromolyn considerably increased microvascular neutrophil crawling velocities in cQD-exposed mice, surpassing even the level of the control group (Figure 4g), most pronounced in the 15-30 min period (Figure 4g). This suggests that cellular degranulation slows down neutrophils in the proximity of deposited NPs, potentially intensifying crawling and probing behavior and subsequent leukocyte recruitment to the airspace in these areas.

To characterize the release of pro-inflammatory mediators 1h after inhalation of cQDs and CNPs and assess the impact of cromolyn treatment, chemokine concentrations in BAL supernatants were quantified by use of a multiplex cytokine assay. After 90 min of nanoparticle exposure, inflammation-related factors in bronchoalveolar lavage (BAL) from mice were analyzed. Both, cQDs and CNP inhalation resulted in a swift release of the neutrophil attractants CCL3, CXCL5 and IL1b. Interestingly, a rapid release of inflammatory factors was not observed in PEG-QDs exposed mice, which is also consistent with the lack of subsequent induction of neutrophilic inflammation after inhalation (Figure 4h and see below Figure 6h, i). Furthermore, the rapid release of inflammatory factors could be effectively inhibited by application of the degranulation inhibitor, cromolyn (Figure 4h). Even if local cytokine changes might be blurred upon BAL analysis, these results suggest that inflammatory factors, including the CXCL family (Figure 4i), are involved in the rapid recruitment of neutrophils and might be released during cellular degranulation following exposure to NPs.

In addition, a focused examination of the BAL levels of the cytokine TNFα after NP inhalation was conducted due to recent findings indicating its crucial role in neutrophil extravasation in skin inflammation by promoting intraluminal crawling^40^. As shown in Figure 4j, significantly increased TNFα levels were detected 2 h after CNP inhalation, but not for cQDs. Furthermore, TNFα neutralization by oropharyngeal aspiration of anti-TNFα mAbs 3 h prior to cQD or CNP inhalation resulted in significantly reduced neutrophil accumulation over the observation period (Figure 4k, l), indicating that TNFα signaling in the airspace is indeed involved in the rapid neutrophil recruitment upon NP inhalation.

### AM patrolling ability being a prerequisite for NP clearance and subsequent neutrophil recruitment

Next, we investigated whether AM patrolling and/or AM phagocytosis of NPs are critical for the spatially and temporally defined immune response. First, we addressed whether AM crawling, which was enhanced after NP inhalation (Fig 3f, g, h) is a prerequisite for the initiation of neutrophil recruitment. For this, we targeted the adhesion molecules CD11a/LFA- 1 which are highly expressed on AMs and has been shown to be utilized by AMs to crawl on alveolar epithelium^26^ and its epithelial expressed adhesion ligand ICAM-1, which is also implicated in leukocyte migration into the lung^40–42^. ICAM-1 and LFA-1 were neutralized in separate experiments by administration of the respective blocking antibodies, by oropharyngeal aspiration 3 h prior to NP-inhalation into the lungs of mice (Fig 5a). AM motility in anti-LFA- 1, as well as anti-ICAM-1 mAb treated mice was significantly decreased following cQDs and CNP inhalation, compared with isotype-antibody treated control mice (Figure 5b-f).

**Figure 5.**
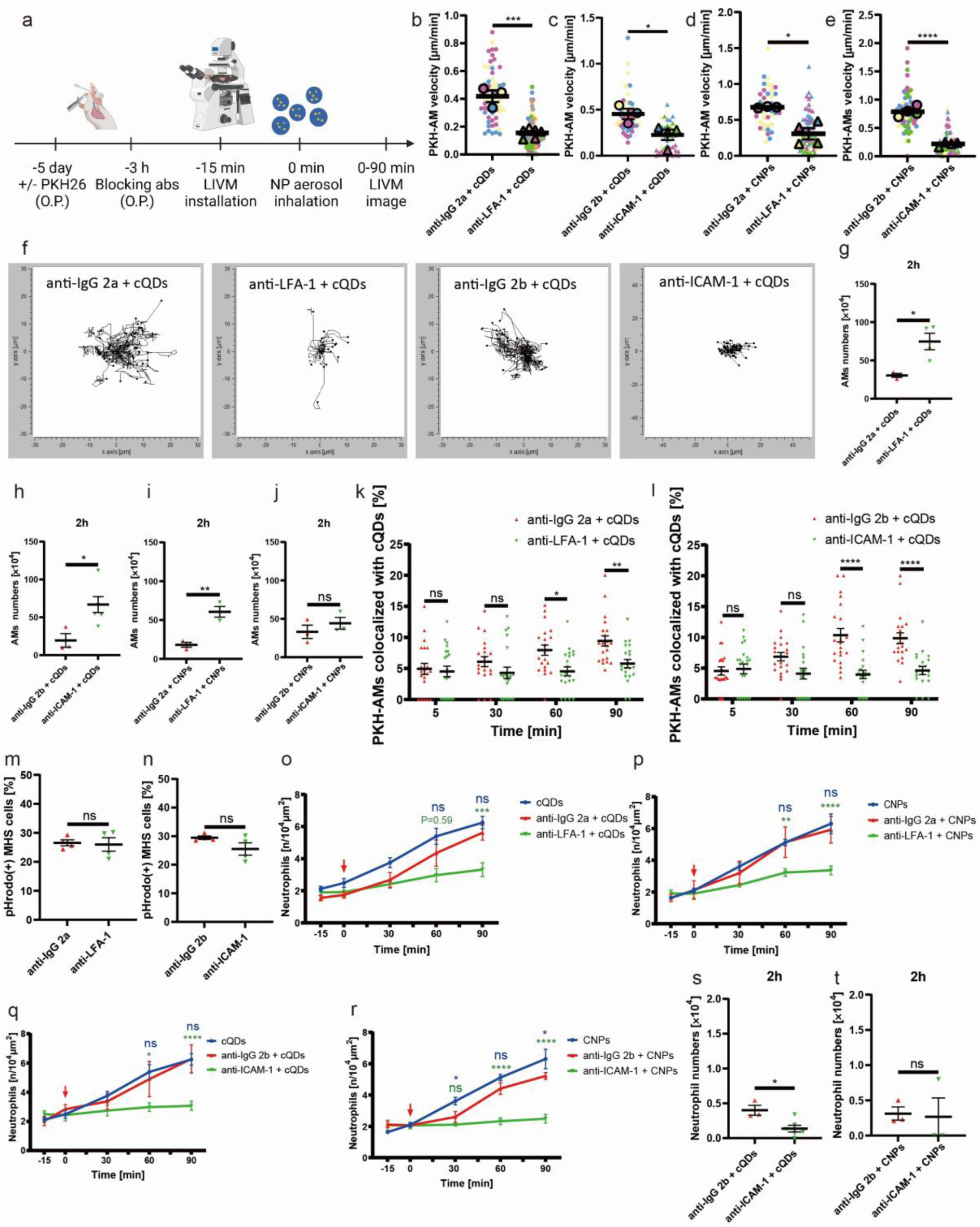
Inhibition of alveolar ICAM-1 and LFA-1 attenuates the uptake of NPs by AMs as well as the subsequent neutrophil recruitment. (a) Schematic of ICAM-1 and LFA-1 neutralizing experiments followed by NPs inhalation in PKH26 pretreated mice. (b–e) Average track velocity (µm/min) of AMs in respective isotype control mAbs, anti-LFA-1, or anti-ICAM-1 mAbs treated mice followed by cQDs or CNP inhalation exposure during a 1 h L-IVM imaging session, n=3-6 mice/group. The mean values of the individual mice are shown as larger circles and triangles of the same color. (f) Spider plot of randomly chosen PKH-AM migration paths in the alveoli of isotype, anti-LFA-1, or anti-ICAM-1 mAbs-treated mice after cQDs or CNPs exposures during a 1 h L-IVM imaging session (0-1h), 10-20 AMs / mice, n=3-6 mice/group. (g)-(j) Quantification of AMs in BAL fluid of mice treated with anti-LFA-1, anti-ICAM-1 or isotype mAbs prior to cQDs (g and h) or CNP (i and j) 2 h after inhalation, n=3-4 mice/group. (k)-(l) Quantification of QDs uptake by AMs in mice pretreated with anti-LFA-1 (g)/ICAM-1 (h) or isotype mAbs after cQDs inhalation, n=3-4 mice/group. (m)-(n) Phagocytic capability was analyzed by flow cytometry of MH-S cell pretreated with anti-LFA-1, anti-ICAM-1, or isotype mAbs 3 h prior to incubation with PH-dependent fluorescent pHrodo E. coli particles (5 μg/ml) for 1 h in vitro. n=4 independent replicates. (o)-(r) Alterations in neutrophil numbers over time under L-IVM were compared between mice pretreated with anti-LFA-1, anti-ICAM-1 or isotype mAbs followed by cQDs (o and q) or CNP (p and r) inhalation, n=3-6 mice/group. (s)-(t) Quantification of neutrophils in BAL fluid of mice treated with anti-ICAM-1 or isotype mAbs prior to cQDs (s) or CNP (t) 2 h after inhalation, n=3-4 mice/group. Data are presented as Means ± SEM. ns: p ≥ 0.05, *: p < 0.05, **: p < 0.01, ***: p < 0.001 and ****: p < 0.0001.

Interestingly, following cQDs as well as CNP inhalation, more AMs could be recovered by lavage from anti-LFA-1 and anti-ICAM-1 antibody-treated mice compared to mice receiving isotype-matched control antibodies (Figure 5g-j). This implies that LFA-1/ICAM-1 blockade also impaired the ability of AMs to firmly anchor to the alveolar walls in vivo, leading to a significant reduction in the adhesion strengthening between AMs and lung epithelial cells, thereby further affecting the crawling and patrolling of AM on alveolar epithelial cells.

Attenuated AM patrolling due to LFA-1/ICAM-1 blockade, affected the clearance of deposited NPs by AMs. Pre-treatment via the airspace with anti-LFA-1 as well as anti-ICAM-1 mAbs prior to cQDs inhalation induced a significant reduction in cQDs internalization in AMs at 60 and 90 min as compared to isotype mAb-treated mice (Figure 5 k, l).

AM phagocytosis capability was seemingly not altered by the application of anti-LFA-1 and anti-ICAM-1 antibodies, since MH-S cells incubated with anti-LFA-1 or anti-ICAM-1 antibodies for 3 h displayed no significant difference in phagocytosis of pHrodo E. coli particles^43^ at 60 min compared with isotype-treated MH-S cells (Figure 5 m, n).

Taken together, this suggests that the lower efficiency of NP phagocytosis by AMs in vivo upon blocking LFA-1/ICAM-1 is due to the mitigating effect on AM motility rather than on AM phagocytosis. This supports the view that AMs rely on ICAM-1/LFA-1 to migrate on the alveolar epithelium to clear deposited NPs in the alveoli.

In addition, prior application of anti-LFA-1, as well as anti-ICAM-1 mAbs, also diminished neutrophil recruitment induced by cQDs and CNP inhalation effectively, as compared to the respective isotype mAb and vehicle controls at 60 and 90 min in L-IVM experiments (Figure 5o, p, q, r). This was partially confirmed by quantification of neutrophils in the BAL fluid of these mice at 2 h (Figure 5s, t).

To rule out that the diminished neutrophil recruitment is caused by the diffusion of antibodies across the epithelial barrier and subsequent direct inhibitory interaction with endothelial cells and neutrophils, we administered LFA-1 and ICAM-1 blocking antibodies in separate experiments, via i.v. injection, directly into the circulatory system at concentrations which blocked TNF-α-induced (for LFA-1)^44^ or cQD-induced (for ICAM-1)^28^ leukocyte recruitment in skeletal muscle tissue. Intravenous application of anti-ICAM-1 blocking mAb in mice receiving cQDs by inhalation, elevated the numbers of recruited neutrophils in the lungs, compared to WT mice receiving cQDs, whereas anti-LFA-1 mAbs attenuated cQD induced neutrophil recruitment, thus indicating an involvement in intravascular neutrophil recruitment (Figure S3a, b). Taken together, the above results support that blocking epithelial ICAM-1 and AM LFA-1 results in the inhibition of cQD and CNP-induced neutrophil recruitment and provide insights into their roles in the processes taking place in the intravascular compartment.

### NP phagocytosis by AMs is a key event in the initiation of rapid neutrophil recruitment

‘Stealth’ QDs which are not readily recognized for phagocytosis by macrophages were applied to further characterize the role of NP phagocytosis by AMs in the initiation of neutrophil recruitment. Their ‘stealth’ feature is due to the modification/coating of the QD surface with polyethylene glycol (PEG), which largely reduces bio-molecule binding^28,45–47^. For this purpose, amino-PEG-QDs (aPEG-QDs) of the same size were used, which show identical deposition efficiency (0.51±0.17%, Figure 6a) and dose as cQDs in the lungs after aerosolization and inhalation (deposited dose determined by quantitative fluorescence: 16 cm^2^/g, Figure 6a).

**Figure 6.**
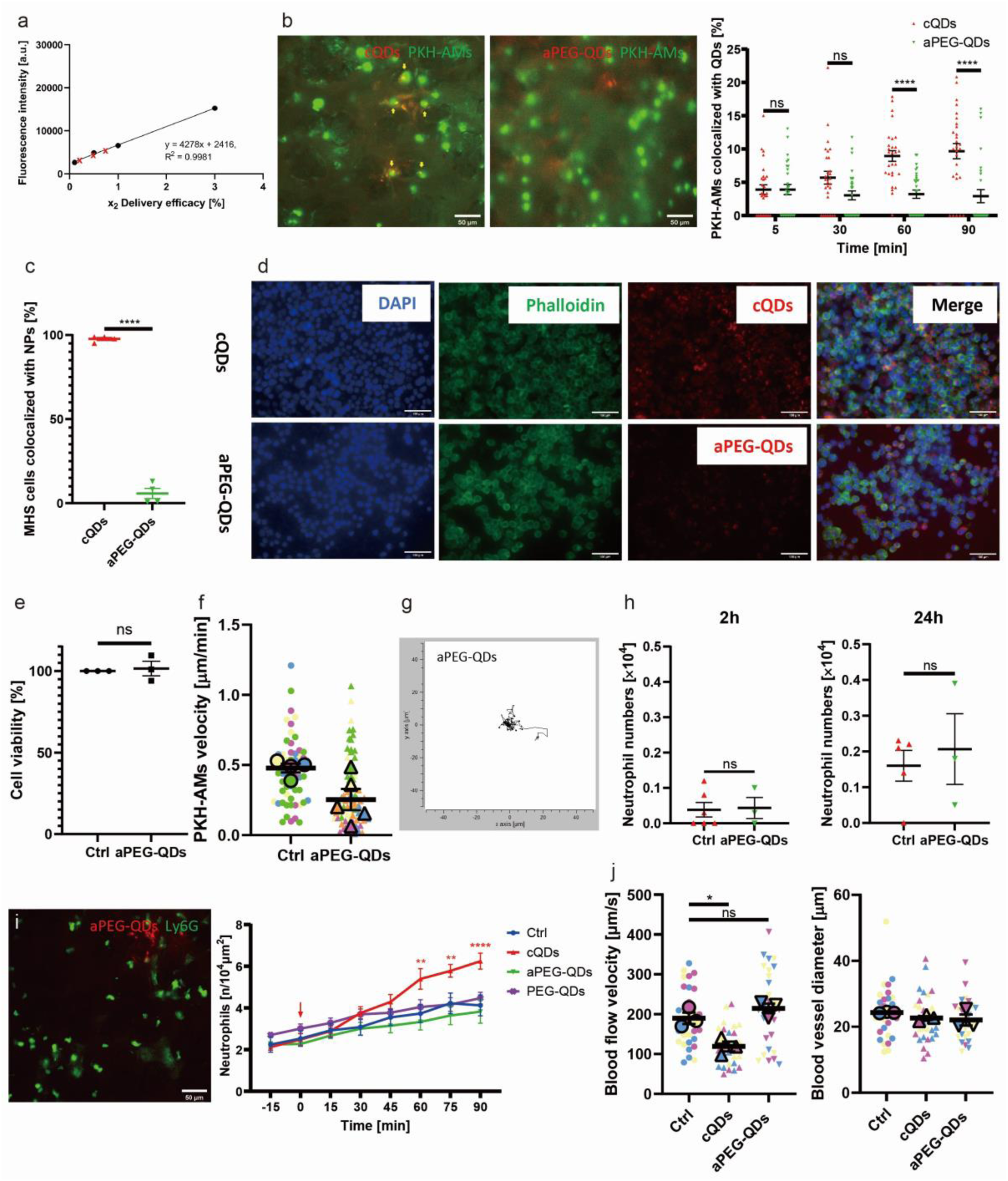
PEGylation reduces NP-induced neutrophil recruitment due to preventing the recognition and internalization of NPs by AMs. (a) Quantification of lung deposited aPEG-QDs after inhalation, calculated according to the linear fluorescence intensity− standard curve (R^2^ > 0.99) obtained from blank lung homogenates spiked with known aPEG-QDs doses (black spheres). Red crosses indicate the respective aPEG-QDs doses (0.51±0.17%, Mean±SD, normalized to the invested dose) detected in mice lung homogenates of 3 independent inhalation applications, n=3. The deposition efficiency and thus the lung-deposited dose (16 cm^2^/g) correspond to that after cQDs inhalation (Figure 1e). (b) L-IVM images at 90 min showed QD (red) dependent uptake (yellow) by PKH-AMs (green), scale bar: 50 μm (left: cQD and aPEG-QD on the left and right, respectively). Quantification of AMs colocalized with QDs, indicating cellular uptake in cQDs-exposed and aPEG-QDs-exposed mice using L-IVM. Here, 7 randomly selected fields of views per time point and mouse were analyzed, n=3 mice/group (right). (c) Quantification of in vitro uptake of cQD and aQD by (alveolar macrophage-like) MH-S cells using FACS analysis 1 h after QDs incubation, n=4. (d) Epi-fluorescence microscope images of cQDs and aPEG-QDs (red) of MH-S cells (phalloidin stained, green) grown at the air-liquid interface 2 h after NP exposure with the VITROCELL Cloud system. DAPI-stained nuclei are depicted in blue. Scale bar: 100 μm. (e) M-HS cells were exposed to aPEG-QDs (8nM) for 1h and cell viability was determined by WST assay. (f) Average track velocity (µm/min) of AMs in different NP-exposed mice (1h) during a 1 h L-IVM imaging session. Each symbol represents an individual random AM combined from 4-5 separate mice. The mean values of the individual mice are shown as larger circles and triangles of the same color. (g) Spider plot of randomly chosen PKH-AM migration paths in the alveoli of mice after different aPEG-QDs exposures during a 1h L-IVM imaging session (0-1h). 10-20 AMs / mouse; n=4-5 mice. (h) Quantification of BAL neutrophil numbers 2 h or 24 h after aPEG-QD and vehicle inhalation, n=3-6 mice/group. (i) Left: L-IVM images at 60 min after aPEG-QDs and PEG-QDs (red) inhalation, anti-Ly6G mAbs labeled neutrophils (green), scale bars: 50 μm. Right: Quantitative L-IVM analysis of neutrophil numbers after cQD, aPEG-QD, PEG-QD and vehicle inhalation. The deposited dose of QDs is 16 cm^2^/g (geom. surface area of NPs / mass-lung), n=6 mice/group. (j) The diameter of alveolar microvessels in the observation area is not affected by QD exposure (left). Quantitative analysis of blood flow velocity (movement of fluorescent beads in blood vessels) 90 min after inhalation of cQDs, aPEG-QDs, PEG-QDs and vehicle (right). The mean values of the individual mice are shown as larger circles and triangles of the same color. Data are presented as Mean ± SEM for n=3 mice/group. ns: p ≥ 0.05, *: p < 0.05, **: p < 0.01, ***: p < 0.001 and ****: p < 0.0001.

Low levels of AMs colocalized with QDs at 5 min after inhalation (median: < 3%) and the percentage of colocalization is gradually increasing for cQDs over the observation time of 90 min, but not for aPEG-QDs as detected by L-IVM (Figure 6b). To investigate this further, MacGreen mice were utilized, where Csf1r-EGFP is expressed selectively in macrophage and monocyte cell lineages. Accordingly, higher numbers of QD-positive AMs were observed in precision cut lung slices obtained from MacGreen mouse lungs having inhaled cQDs, this is not seen for aPEG-QDs (Figure S4a, b).

To verify the QD-dependent uptake by AMs in vitro, we exposed alveolar macrophage-like MH-S cells, cultivated at the air-liquid interface^48^, to aerosolized cQDs or aPEG-QDs for 1h. As quantified by FACS analysis and confirmed by confocal microscopy of exposed MH-S cell cultures, 95.8±1.8 % MH-S cells internalized cQDs nanoparticles, while only 8.8±1.5 % MH- S phagocytosed aPEG-QDs (Figure 6c, d). Cell viability of MH-S cells was not reduced by aPEG-QDs, i.e. the low uptake of aPEG-QDs is not due to cytotoxicity (Figure 6e)

Returning to in vivo conditions, L-IVM revealed that the average AM patrolling velocity increases in the presence of cQDs relative to the negative control group, while it decreases after inhalation of aPEG-QDs (Figure 6f, g).

To substantiate AM phagocytosis as the key event for NP inhalation triggered neutrophil recruitment, we additionally used pure PEG-QDs, without amine surface functionalization. All three types of QDs exhibit the same core structure, display identical diameter (20 nm) following nebulization from suspensions (Figure S5a-c), and appear uniformly distributed through the whole lungs after inhalation (Figure S5d-f), with comparable lung delivery efficiency, as assessed by fluorescence imaging in excised lungs (Figure 1e, 6a). Merged images of maximum intensity projections (MIP) of QD fluorescence signals with the 3D lung structure confirm that cQDs as well as aPEG-QDs exhibited a uniform distribution in the lungs in the overall view (Figure S5j-k). However, some local differences existed at the microscopic scale. After cQDs exposure, preferential central acini (blue arrow) exhibited strong cQD fluorescence while in the great majority of the peripheral acini (yellow arrow), fewer cQDs were deposited (Figure S5j, l). Conversely, after aPEG-QDs inhalation, the distribution of aPEG-QDs was similar in central or peripheral areas of the lungs (Figure S5k, m). L-IVM revealed that inhalation of cQDs and PEG-QDs resulted in a roughly 2-fold lower average fluorescence intensity in the respective fields of view than aPEG-QD inhalation (Figure S5g-i and Figure S5n).

In contrast to cQDs, inhalation of aPEG-QDs and PEG-QDs did not result in rapid neutrophil recruitment detected by L-IVM relative to vehicle control after QD inhalation (Figure 6i). This lack of aPEG-QD induced neutrophil recruitment is corroborated by unchanged neutrophil numbers in BAL at 2 h and 24 h after aPEG-QDs inhalation (Figure 6h). Due to the lack of an inflammatory response to these “stealth” QDs, also blood flow velocities remained unchanged after inhalation of both aPEG-QDs and PEG-QDs (Figure 6j).

To investigate cellular uptake and subcellular distribution of NPs, transmission electron microscopy was applied to the lungs of mice after exposure to cQDs, aPEG-QDs or PEG-QDs for 1 hour. Some cQD clusters were identified within the cytoplasm or in endolysosomes of AMs, whereas no such aggregated QDs could be identified in AMs in the PEG-QD or aPEG- QD groups (Figure S6a). Interestingly, cQDs were found to be attached to or agglomerated with alveolar surfactant (tubular myelin sheets, based on their ultrastructure), at the air-liquid epithelial interface, which was not observed for aPEG-QDs and PEG-QDs exposed mice (Figure S6b). The TEM results confirm the lack of AM uptake for aPEG-QDs and PEG-QDs and thus support the stringent necessity of NP phagocytosis by AMs for the initiation of the inflammatory response.

### C5aR1 and FcγRI functions are crucial for rapid neutrophil recruitment upon NPs exposure

Since NP phagocytosis was identified as a key event, we were interested in the involved uptake receptors. The complement receptor C5aR1 (CD88) is highly expressed on AMs^26,49^ (Figure 7a), and thus served as a good candidate. AMs are known to get rapidly stimulated by C5a to chemotaxis and uptake of bacteria^26^ and C5a is closely related to cellular degranulation^50,51^. To assess if the C5a-C5aR1 axis was vital in neutrophil recruitment after NP inhalation, we applied C5aR1-blocking (CD88) mAbs by oropharyngeal aspiration, followed by cQD and CNP aerosol inhalation. Indeed, the application of anti-C5aR1 mABs effectively prevented the rapid neutrophil recruitment after cQD-inhalation (Figure 7c), as well as after CNP-inhalation (Figure 7d). Likewise, only baseline levels of cQD uptake by AMs were detected in the intravital images of anti-C5aR1-treated mice as compared to isotype mAb-pretreated mice (Figure 7e), and, upon cQDs exposure, AM velocity was not affected by anti-C5aR1 mAbs pretreatment (Figure 7g).

**Figure 7.**
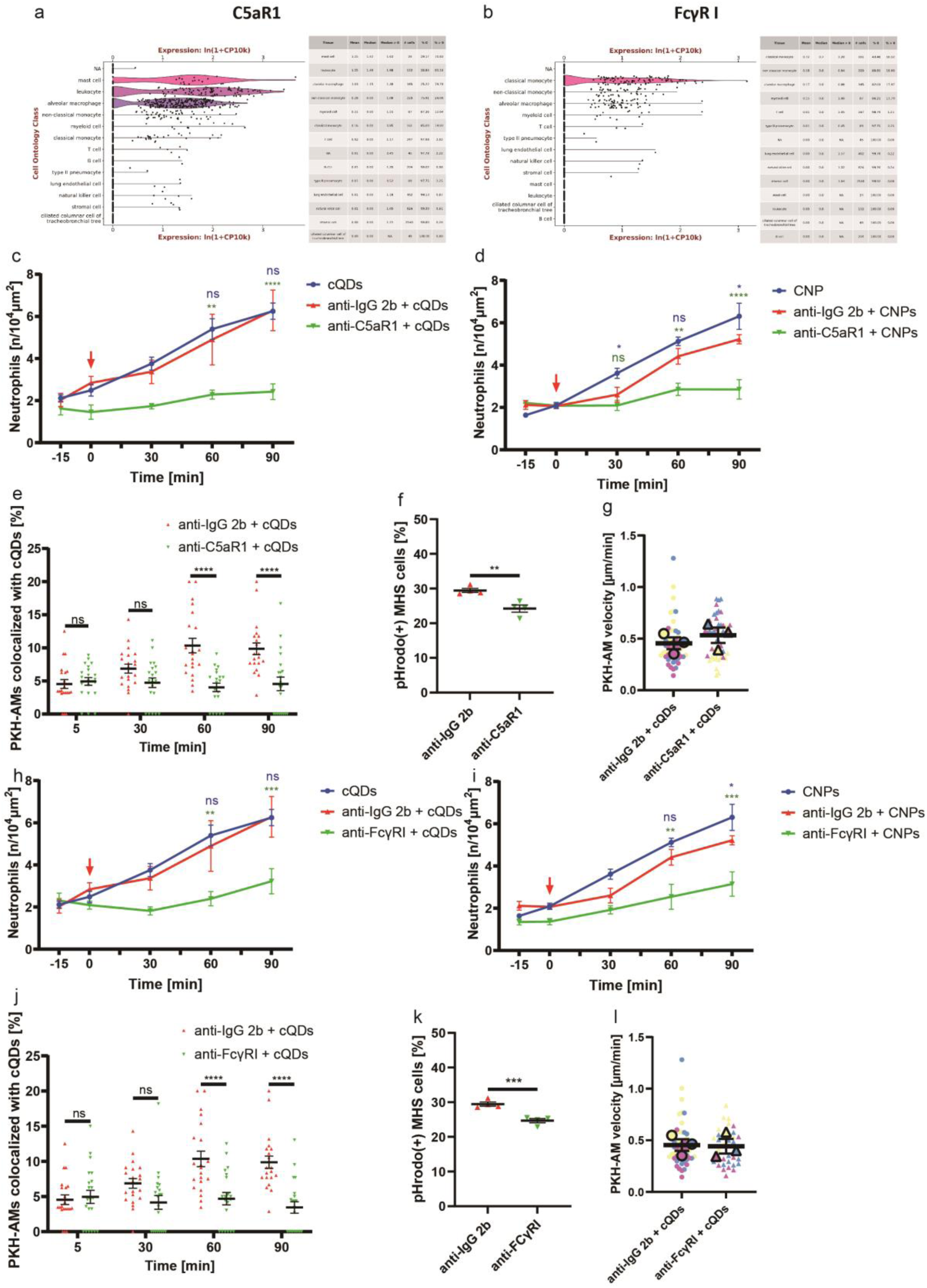
AM FcγRI and C5aR1 are involved in the rapid neutrophil recruitment upon NP exposure. (a)-(b) C5aR1 and FcγRI expression in the lungs of naive mice. C5ar1 and FcγRI are highly expressed on the surface of AMs and monocytes, but not on alveolar epithelial cells^49^. (c)-(d) The time course of neutrophil levels according to L-IVM was compared between mice pretreated with anti-C5ar1-mAb or isotype mAb before inhalation of cQDs (c) or CNPs (d) or vehicle-control, n=3-6 mice/group. (e) Quantification of cQDs uptake by AMs in mice pretreated with anti-C5aR1 mAbs or isotype mAbs 3 h before cQDs inhalation, n=3 mice/group. (f) Phagocytic ability of MH-S cells treated with anti-C5aR1-mAbs or isotype control mAbs 3 h prior to pHrodo E. coli (5 μg/ml) incubation for 1h in vitro was analyzed by flow cytometry. n=4 times/group. (g) Average track velocity (µm/min) of AMs in isotype or anti-C5ar1 mAbs treated mice followed by cQDs inhalation during a 1h L-IVM imaging session. The mean values of the individual mice are shown as larger circles and triangles of the same color, n=3 mice/group. (h)-(i) Alterations in the amounts of neutrophils over time under L-IVM were compared between the mice pretreated with anti- FcγRI or isotype mAbs before inhalation of cQDs (h) or CNPs (i) or vehicle-control, n=3-6 mice/group. (j) Quantification of QDs uptake by AMs in mice pretreated with anti-FcγRI mAb or isotype mAb 3 h before cQDs inhalation, n=3 mice/group. (k) In vitro AM phagocytosis of pHrodo E. coli was analyzed by flow cytometry using AM-like MH-S cells treated with anti- FcγRI mAb or isotype mAb control 3 h before pHrodo E. coli (5 μg/ml) application (1h), n=4 times/group. (l) Average track velocity (µm/min) of (in vivo) AMs in isotype or anti-FcγRI mAb-treated mice followed by cQDs inhalation exposure during a 1h L-IVM imaging session. The mean values of the individual mice are shown as larger circles and triangles of the same color, n=3 mice/group. Data are presented as Mean ± SEM. ns: p ≥ 0.05, *: p < 0.05, **: p < 0.01, ***: p < 0.001 and ****: p < 0.0001.

Investigating pathogen phagocytosis using pHrodo E-coli particles exposed to AM-like MH-S cells for 3 h with anti-C5aR1 mAbs under in vitro conditions, showed only slightly decreased uptake at 60 min (Figure 7f), which suggests different receptors for pathogens and sterile particle.

Taken together, these results suggest that blocking C5aR prevents NP-induced neutrophil recruitment due to reduced phagocytic ability/recognition of NPs by AMs without affecting AM patrolling behavior. Like F4/80 or the protein tyrosine kinase MER (MERTK), FcγR I is specifically expressed on the surface of mouse macrophages^52^ (Figure 7b). Based on earlier studies in primary murine macrophages, FcγR knock-out mouse models, and human monocyte- derived macrophages, FcγR-mediated phagocytosis was more efficient at inducing inflammation than CR-mediated phagocytosis^53,54^. Since also FcγR is closely involved in the degranulation of many kinds of inflammatory cells, such as NK cells and mast cells^55–58^, it presented another candidate uptake receptor to be investigated. To determine FcγR I function on NP-induced neutrophil recruitment, we blocked FcγR I with anti-CD64 mAbs administered into the lungs 3 h before NPs inhalation. 60 min after cQDs or CNP exposure, neutrophil accumulation in the lungs of anti-CD64 treated group was reduced compared to the isotype control group (Figure 7h, i). After anti-FcγR I mAbs treatment, NPs phagocytosis of AMs was again greatly reduced compared to isotype mAbs treated mice (Figure 7j), while it had no effect on the patrolling movement of AMs (Figure 7l). Again pHrodo E. coli phagocytosis by MH-S cells was only slightly attenuated by incubation with CD64 blocking mAbs, in comparison to the isotype-treated group (Figure 7k). Taken together, these results support NP phagocytosis by AMs as a crucial event in the NP-induced alveolar neutrophil recruitment cascade, presumably followed by the release of proinflammatory mediators by AMs.

## Discussion

Exposure to nanoparticles, especially through ultrafine particulate matter, a crucial component of ambient air pollution, is unavoidable in everyday life and is associated with significant health risks, particularly concerning the development of cardiovascular diseases^59^, but also in terms of acute and chronic respiratory symptoms and diseases^60–62^. Nano-sized particles specifically have been shown to be of great concern as they effectively reach the alveolar space and have a larger mass-specific surface area, which is one of the main drivers of particle-induced lung inflammation^63^. In addition to a possible translocation of nanosized particles, the immediate inflammatory reaction in the lungs is considered to be critical for cardiovascular consequences^8,64^. However, the initiation of complex innate immune reactions, which is triggered by NPs in humans as well as in mice, remained largely unresolved so far. While it is widely accepted that particle surface interactions with biological membranes trigger oxidative stress ^65^, the specific cellular players responsible for triggering neutrophil recruitment at nanoparticle deposition sites are still unclear. Prime candidates are tissue-resident AMs, previously called ‘dust cells’, which as resident phagocytes of the alveoli, play an essential role in the homeostatic “vacuum-cleaning” of pulmonary surfactant turnover, daily cellular debris and inhaled dust with immunosuppressive function to maintain non-inflammatory conditions^66^. AMs are essential for pathogen elimination via non-specific phagocytosis and are strategically placed to initiate a robust inflammatory response to more pathogenic stimuli^19^. Accordingly, AMs are also involved in the clearance of inhaled nanomaterials^12^, but their role in the initiation of inflammation upon pulmonary deposition of sterile, pyrogen-free particles is controversial. Depending on the kind and cytotoxicity of inhaled particulate matter, AMs have been considered to become activated upon contact or uptake and to produce pro-inflammatory cytokines, or even undergo various cell death pathways. However, these findings are usually based on in vitro studies obtained for ambient particulate matter samples containing various amounts of endotoxin or very high doses of highly toxic materials (e.g. residual oil fly ash of 100 µg/ml medium, i.e. nominal cellular dose of 0.3 µg/cm^2^ cells corresponding to 114 years of realistic urban particle exposure at 10 µg/m^3^)^67–69^. In contrast to the common assumption of a pro-inflammatory AM activation by inhaled soot-like nanoparticles, our earlier in vivo study showed at least no transcriptional activation of AMs in the lung of mice 6 and 12 h after intratracheal instillation of 20 µg CNPs^7^. In this study we used different spherical (10-20 nm) nanoparticles to explore the initiation of lung inflammation, employing fluorescent QDs with two different surface modifications, for tracking particle deposition and uptake, and soot-like CNPs.

The accumulation of neutrophils as the first responders of the innate immune system is considered the hallmark of inflammation^21^. Also, upon nanoparticle exposure neutrophils are rapidly recruited to the airspace, as described in various rodent models^7,10^. Both AMs and neutrophils are sentinel cells that patrol, explore, and eliminate threats within the host^21,26^. To date, the functions of these two types of immune cells during short-term exposure to inhaled NPs remain poorly understood and their interplay in the response to alveolar deposited NPs has not been investigated in their natural environment, i.e. the breathing lung. Combining lung intravital microscopy and ventilator-assisted NP inhalation, enabled us for the first time to study dose-aware NP deposition, AM motility and cellular NP uptake in the living organism, and therefore to identify key events leading to spatiotemporally resolved neutrophil recruitment induced by inhaled NPs.

Utilizing the PKH-26L labeling method for resident phagocytes implemented by the Kubes laboratory to visualize AMs, our study confirmed their finding that on average one AM maintains the homeostasis of three murine alveoli^26^. AMs exhibited crawling movements necessary to clean the alveolar surface from aged and excessive surfactant, cell debris and NPs. Intriguingly, in our L-IVM study, inhalation of both cQD and CNP at equivalent inflammogenic dose levels almost doubled the AM migration velocity from around 0.5 µm/min up to 1 µm/min, an effect not observed for PEG-shielded aQDs. Notably, while AM migration patterns in the alveoli appeared to be randomly oriented, directed movement toward NP clusters was observed over short distances (approx. 10-20 µm). The observed net increase of AM numbers in the close vicinity of deposited NP ‘hotspot’ might have been facilitated by an accelerated random walk. The change in AM migration thereby represents the first cellular event, initiating the pulmonary inflammatory response, as spatial mapping clearly indicated that NP-internalizing AMs prompted neutrophils to preferentially arrest in nearby microvessels.

The crucial role of AMs in pulmonary inflammation has been well-documented, particularly in models of fiber-induced inflammation by experimental depletion of AMs, which decreased carbon nanotube induced neutrophilic inflammation ^70^. In our hands, using established protocols for clodronate liposome mediated AM depletion resulted in elevated neutrophil numbers in the airspace, higher than those induced by the here used low dose NP inhalation challenge, rendering this approach at least for our sensitive experimental approach useless. In addition, recent critiques of clodronate liposome methodologies raise concerns about the interpretation of such studies ^71^.

Effective nanoparticle clearance by AMs hinges on both their motility and phagocytic capabilities. Our data indicated that AMs react to the alveolar presence of NPs, (CNP and cQDs) with increased motility, which turned out to be dependent on the expression of ICAM- 1, the adhesion ligand for the β2 integrin LFA-1. ICAM-1 is highly expressed by murine type 1 pneumocytes, is upregulated in inflammatory lung disease^72^. LFA-1 (*Itgal*) gene knockout weakened AMs motility and impaired clearance of endogenous cellular debris in the alveolar space^26^. In our study, both anti-LFA-1 and anti-ICAM-1 pre-treatment alleviated the cQD- induced increase in AM patrolling speed. Importantly, LFA-1 or ICAM-1 blockade did not affect the phagocytic ability of AMs in *in vitro* experiments. Therefore, inhibition of LFA-1 and ICAM-1 is expected to reduce clearance of cQDs by AMs due to reduced AM motility in the alveoli, but not due to mitigated AM phagocytic capacity. AM-epithelial interactions were also required to trigger early neutrophil attraction, as LFA-1 and ICAM-1 blocking antibody application eliminated the rapid recruitment of neutrophils occurring within the first hour after NP inhalation.

The alveolar surfactant layer plays a pivotal role in preventing alveolar collapse by reducing surface tension, immunologically serving as functions the first biological barrier against inhaled threats such as NPs. With its protein-rich lipids, lung surfactant can generate a bio- corona wrapping alveolar deposited NPs, subsequently modulating NP-cell interactions^73,74^. Our transmission electron microscopy (TEM) images showed the association of cQDs with alveolar surfactant structures highlighting the high affinity of cQD-NPs for biomolecular interactions. AMs possess various surface receptors for pathogen identification and internalization, ^75,76^, with the particle uptake potentially facilitated by opsonization via complement factors or surfactant proteins. In this study, targeting FcγRI and C5aR1, both highly expressed on the surface of AMs but not on lung epithelial cells, eliminated NPs-induced neutrophil recruitment without altering the patrolling status of AMs and resulted in defective NP recognition/phagocytosis by AMs. The surfactant abundant in the alveoli is primarily the complex secreted by alveolar type II epithelial (ATII) cells and their recycling and clearance are completed by the cooperation of ATII and AMs^77^. PEGylation forms a steric barrier to reduce the opsonization of NPs^45^, allowing PEGylated QDs to evade phagocyte recognition ^28^. In contrast, non-PEGylated cQDs formed a biocorona, enhancing their uptake by AMs. Indeed, the uptake of PEGylated QDs by AMs and the subsequent recruitment of neutrophils was significantly reduced, indicating that the surface area of the lung deposited NPs, which proved to be the most reliable dose metric for predicting acute NP lung toxicity^5^, reflects the NP availability for cellular (AM) uptake.

Various immune cells including NK cells, eosinophils, and mast cells, store inflammatory mediators in intracellular vesicles known as granules^78^. These vesicles play a crucial role in releasing mediators during the host immune response to ensure immediate cell communication and defense. Previously, our group demonstrated that preventing cellular degranulation with cromolyn completely eliminated cQDs-evoked leukocyte recruitment in skeletal muscle postcapillary venules^28^. Consistent with these findings, Lê et al. revealed cromolyn pretreatment greatly reduced TNF-α, IL-1β levels as well as lung MPO activity in LPS-treated mice ^79^. A direct involvement of alveolar-localized mast cells in local neutrophil recruitment seems actually less likely, since mast cells are predominantly located in the central airways rather than peripheral alveoli of mice^80^, which is in accordance with our own results, in which we could not detect mast cells in the alveolar regions, where L-IVM analysis was conducted^81^.

Moreover, the presence of mast cells in the lung parenchyma has recently been linked to the hygiene status of the mice. Mice housed under ’specific pathogen-free’ conditions, are almost devoid of these cells, in contrast to wild mice^82^. In our study, inhibiting degranulation by cromolyn led to a marked reduction in cQD- and CNP-induced cytokine levels in the BAL to control levels (e.g. CXCL10, -12, CCL3, and IL-1b), and prevented neutrophil recruitment up to 24 h after NP inhalation. These results might provide a new direction for mitigating the harm of NP exposure in the future. Furthermore, our data revealed that AMs served as critical effector cells in NP-induced neutrophil recruitment, a key event also requiring rapid TNFα release. The role of AMs in driving both acute and long-lasting inflammatory responses, after exposure to differently shaped NPs has been further addressed through single cell transcriptomics by our group^83^.

Recently, the function of neutrophils in NP clearance from the bloodstream, before being diverted to the liver, has been hightligted^84^. While the mononuclear phagocytic system primarily handles NP elimination from the bloodstream^85,86^, our observations indicate that neutrophils play a significant role in NP clearance from the alveolar space. With L-IVM, we identified a substantial NP internalization (cQDs) by alveolar localized neutrophils, which were recoverable in BAL. It is however not clear to which extent neutrophil uptake can be generalized for different types of NPs. Notably, there is limited research on neutrophils scavenging inhaled NPs, with most studies focusing on their role in clearing pathogens in lung microbe-infection models^26,87–89^. The involvement of recruited cells such as neutrophils and lymphocytes is understudied in this scenario.

Our L-IVM study revealed tissue-resident AMs are pivotal in initiating the inflammatory immune response to inhaled NPs. AM exposure to deposited NPs increased their crawling speed and accumulation at sites of pronounced particle deposition. Previous studies have shown that epithelial deposition of inhaled NPs is not homogeneous but NPs rather deposit at hotspots within the acinus, the alveolar sacculi, distal to the bronchial-alveolar duct in mice^90^. Whether AMs get attracted to deposition sites or accumulate there upon random migration remains still unclear. However, we show that AM-mediated NP clearance by phagocytosis is a key event required for neutrophil recruitment. Accordingly, neutrophil extravasation is localized to NP deposition hotspots and AM presence as well. Upon particle exposure, neutrophil chemoattractant such as CXCL-1, -2, and -5 and TNFα are rapidly released into the airspace^7^, although whether they initiate leukocyte recruitment or merely amplify the ongoing response has so far not been shown. In our study, we identified cellular degranulation and TNFα signaling as key factors, required for airspace neutrophilia implying that TNFα might be released from NP-phagocytosing AMs. However, at the in vitro level CNP exposed AMs remained transcriptionally quiescent^91^. Particle phagocytosis, involving NP recognition via FcγRI and C5aR1, nevertheless was essential to initiate lung inflammation, further underscoring the role of AMs as key cells eliciting the response to inert NPs, like CNPs.

## Conclusion

In summary, our study demonstrated that alveolar-deposited NPs prompt a rapid inflammatory response that is initiated by intensified AM patrolling behavior and particle phagocytosis. Notably, it was found that the early neutrophil responses are locally focused to those alveoli, which have been challenged by a high burden of deposited NPs. However, only NPs, which could be recognized and phagocytosed by AMs induced an immune response, whereas “stealth” NPs, such as PEGylated QDs did not. Our data identified NP uptake by AMs as the key event for the initiation of the pulmonary inflammatory response and shed new light on mediators and receptors involved in the initiation of NP-induced sterile inflammation. These new insights allow us to propose novel strategies to regulate AM activity, thus opening potential new therapeutic approaches ranging from air pollution and nanoparticle exposure to chronic inflammatory lung disease.

## Materials and Methods

### Mice

C57BL/6 (WT) mice were purchased from Charles River (Sulzfeld, Germany) for the respective experiments and Macgreen (Csf1r-EGFP) mice were originally purchased from the Jackson Laboratory (Bar Harbor, ME, USA) and bred in-house. All mice were housed in individually ventilated cages supplied with filtered air in a 12 h light/12 h dark cycle within a specific double-barrier, pathogen-free unit at Helmholtz Zentrum München. Mice were fed with autoclaved rodent feed and water ad libitum. All experiments were performed with female animals at 10 to 16 weeks of age. All protocols used were according to the guidelines drafted by the Regierung von Oberbayern (District Government of Upper Bavaria).

### Nanoparticles

Quantum Dots (QDs): Qdot™ 655 ITK™ Carboxy (cQD), Qdot™ 655 ITK™ Amino (PEG) (aPEG-QDs) and Qtracker™ (PEG) 655 quantum dots (PEG-QDs) were purchased from Invitrogen Corporation (Karlsruhe, Germany) as 8 μM solution, or 2 µM in case of PEG-QDs with an emission wavelength of 655nm. These QDs are composed of a CdSe core encapsulated with a ZnS shell and an extra layer of polyethylene glycol (PEG-QDs) and with additional amine residues (aPEG-QDs) or without any coating (cQDs). The PEG coating comprises short oligomers (1.3 kDa)^46^. The core-shell dimensions of the elongated 655-QDs were 10-12 nm. The physical characterization of QDs, including biomolecule surface association was previously performed in our laboratory and has been published^28,46,47^.

For bio-persistent particles, (BET or geometric) surface area per mass of lung has been established as an allometrically scalable, biologically relevant dose metric for pulmonary inflammation murine lung weight: 0.18 g)^5,92^. For conversion of the typically reported molarity of QD suspensions to this unit one needs to consider the molar mass (1.5-2*10^6^ g/mol), primary geometric particle diameter (20 nm), yielding a mass-specific surface area of (16 cm^2^/g-lung). Printex90 carbon black (soot with low organic content) nanoparticles (CNP) were purchased from Degussa (Frankfurt, Germany) (diameter: 14 nm; organic content: 1%; mass-specific (BET) surface area: 272 m^2^/g as described^93^, and prepared in a 5 µg/µl stock suspension (in water).

### In vivo labeling of AMs and neutrophils

The method was performed according to a previously published procedure^26^. PKH26 phagocytic cell labeling kit (Sigma, Burlington, MA, USA) was purchased from Merck KGaA (Darmstadt, Germany). 0.1 ml PKH dye stock (10^-3^ M) mixed thoroughly with 0.9 ml alcohol to make a 100 μM working solution. The working solution was then diluted in Diluent B to prepare 0.5 μM PKH26PCL, serving as the labeling solution for all L-IVM experiments. By oropharyngeal application, mice were given 75 μl of working solution directly into the airway at least 5 days prior to the experiment.

For in vivo labeling of neutrophils, anesthetized mice (MMF i.p.) were administered with 3 µg of fluorescent-labeled anti-mouse Ly6G antibody (Alexa Fluor® 488, clone: 1A8, Biolegend, California, USA) fluorescent-labeled antibodies via i.v. injection at least 30 min prior to NPs inhalation.

### In vivo blocking/inhibiting experiments

Mice were administered blocking antibodies/isotype control antibodies or inhibitors (see Table 1) via the oropharyngeal method, as described above, in the airways. 3 h after administering blocking antibodies, mice were imaged with L-IVM.

**Table 1:** List of blocking antibodies or inhibitors applied into airways.

| Blocking antibody/Inhibitor | Clone | Manufacturer |
| --- | --- | --- |
| **In vivo mAb anti-mouse LFA-1α** | M17/4 | Bio X Cell,<br>Lebanon, USA |
| **Purified anti-mouse CD54 (ICAM-1) Antibody** | YN1/1.7.4 | ATCC,<br>Manassas, USA |
| **Ultra-LEAF™ Purified anti-mouse CD88 (C5aR) mAb** | 20/70 | BioLegend,<br>Fell, Germany |
| **Ultra-LEAF™ Purified anti-mouse CD64 (FcγRI) mAb** | W18349F | BioLegend,<br>Fell, Germany |
| **In vivo mAb rat IgG2a isotype control** | 2A3 | Bio X Cell,<br>Lebanon, USA |
| **Ultra-LEAF™ Purified Rat IgG2b, $\kappa$ Isotype Ctrl antibody** | RTK4530 | BioLegend,<br>Fell, Germany |
| **Purified rat IgG1, $\kappa$ isotype control antibody** | RTK2071 | BioLegend,<br>Fell, Germany |
| **Purified anti-mouse TNF-α antibody** | MP6-XT22 | BioLegend,<br>Fell, Germany |

Mice were pretreated with blocking antibodies/isotype control antibodies or stabilizers (Table 2) applied into the blood circulation via i.v. injection. 30 min after administering blocking antibodies, mice were exposed to NPs.

**Table 2:**
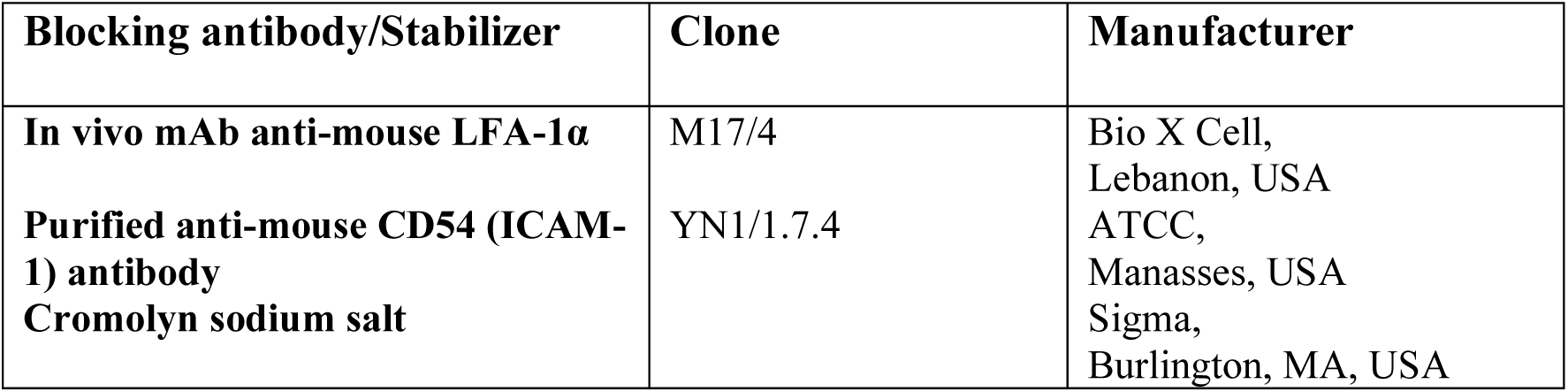
List of blocking antibodies or stabilizers applied into the blood circulation.

### Lung intravital microscopy (L-IVM)

The lung intravital microscope, based on the VisiScope A1 imaging system (Visitron Systems GmbH, Puchheim, Germany), is equipped with an LED light source for fluorescence epi- illumination (CoolLed p-4000, Andover, UK). QDs were excited using the 385nm LED module, melamine resin particles for blood flow tracing were illuminated with the 655 nm LED module, anti-Ly6G-488 was exited with 470 nm, and PKH-26 with 550 nm (all at 50% output power, exposure time 50 ms). The light was directed onto the sample via a quad-band filter (F66-014, DAPI/FITC/Cy3/Cy5 Quad LED ET Set; AHF Analysentechnik AG, Tuebingen, Germany). Microscope images were acquired by a water dipping objective (20x, NA 1.0. Zeiss MicroImaging GmbH, Jena, Germany). A beam splitter (T 580 lpxxr Chroma Technology Corp, Bellows Falls, USA) was used to divide the light from the sample. Images were acquired with two Rolera EM2 cameras and VisiView Imaging software (Visitron Systems GmbH, Puchheim, Germany). The experimental approach was adapted from previously described work^27^. Mice, deeply anesthetized with a mixture of medetomidine (0.5 mg/kg body weight), midazolam (5 mg/kg body weight), and fentanyl (0.05 mg/kg body weight) (MMF) by intraperitoneal injection, and were maintained at 37°C body temperature and monitored using the small animal physiological monitoring (Harvard Apparatus, Massachusetts, United States). Surgical sites (neck and left chest) were locally anesthetized with Bucaine (50 µg/site, Puren Pharma, Germany). Tracheostomy was performed, and a small blunt catheter (20G, B. Braun, Melsungen, Germany) was threaded <5 mm into the trachea, and connected to a small rodent ventilator (MiniVent, Harvard Apparatus, Massachusetts, US). Mice were ventilated with a stroke volume (tidal volume) of 10µl/g body weight (BW) and 150 breaths per minute under 0.1cm/g-BW positive end-expiratory pressure (PEEP) with 100% oxygen.

The mice were then placed in the right lateral decubitus position and a custom made flanged thoracic suction window was inserted into a 5 mm intercostal incision through the parietal pleura between ribs 3 and 4 of the left chest. 20–25 mmHg of suction was used to immobilize the lung by a custom-made system consisting of a differential pressure gauge (Magnehelic, Dwyer Instruments, Inc, USA) and a negative pressure pump (Nupro, St Willoughby, USA). Anesthesia was maintained by the administration of half the first dose of MMF (i.p.) every 45 min.

### NPs aerosol inhalation

For dose-controlled aerosol inhalation, the respiratory parameters of the mechanically ventilated mouse were set to 15 μl/g and 150 breaths/min (miniVent, model 845, Harvard Apparatus, USA). Subsequently, 20 µl of 4 µM (= 7 µg/µl) quantum dot suspension (1:2 diluted stock suspension; diluent: distilled water with 1% saline) or 5 µg/µl CNPs (stock) suspension were nebulized with a breath-activated vibrating mesh nebulizer (Aeroneb Lab Small, volume-weighted droplet diameter (VAD) = 2.5-4.0 μm; manufacturer information VMD Aerogen Inc., Ireland). For this, an infrared-based motion detector was installed in front of the moving ventilator piston for triggering nebulizer activation for 20 ms per breath at the onset of the inhalation phase. After the NP inhalation procedure which typically lasted 1 min for QDs and 5 min for CNPs, the tidal volume was set back to normal breathing conditions (10μl/g BW, 150 breath/min) as used for L-IVM.

### Measurement of blood flow velocity

Melamine resin fluorescent particles (microParticles, Berlin, Germany) (50 μl of 0.05 % stock solution) were injected (i.v.) into mice and imaged at approximately 2.5 frames per second using the fast acquisition mode. The pulmonary blood flow velocity of mice was represented by the trajectory velocity of MF fluorescence particles. The “Manual Tracking” function of ImageJ (National Institutes of Health, Bethesda, USA) was applied to give a specific coordinate axis to the tracing bead location at each time point. Then, bead velocity was analyzed with the

Chemotaxis and Migration Tool Software (ibidi GmbH, Gräfelfing, Germany).

### Quantification of neutrophil kinetics

To quantify neutrophil numbers in the pulmonary microcirculation and the alveolar region, in vivo images of seven observation areas were randomly recorded at intervals of 15 or 30 min pre- and post-inhalation. The L-IVM images were analyzed with Image J software (National Institutes of Health, Bethesda, USA) to quantify the neutrophil numbers. For neutrophil counting, the “Trackmate” function of Image J was used. All neutrophils were automatically counted by the software using the -15 min (neutrophil baseline) image for the threshold and size setting. The same settings were used for subsequent time points, and the results underwent manual calibration.

### BAL preparation and cell differentiation

Mice were anesthetized via i.p. injection of xylazine (5.5 μg / g body weight, WDT, Garbsen, Germany) and ketamine (0.5 mg / g body weight, PharmaWiki, Disentis, Switzerland) and sacrificed by abdominal aorta exsanguination. Immediately after, bronchial alveolar lavage (BAL) was performed by trachea intubation with a 20G cannula (Braun, Melsungen, Germany) and by infusing the lungs 8 times with 1.0 ml sterile phosphate-buffered saline (PBS). The total recovered volume was around 8 ml/mouse. Cell pellets obtained after centrifugation (400 g, 20 min at 4℃) were resuspended in 1 ml sterile PBS and the cell amount was counted using the trypan blue exclusion method. BAL cell differentials were accomplished on cytospin slides with May-Grünwald- Giemsa staining (3 × 200 cells counted).

### Ex vivo 3D light sheet fluorescence imaging of the lung

C57/Bl6 or Csf1r-EGFP mice were sacrificed by exsanguination and transcranial perfused with 30 ml sterile PBS at room temperature to flush all blood (EGFP-labeled cells included) from the lungs. After that, only EGFP-expressing AMs and interstitial macrophages are left in the lung. The lung tissue was stained and cleared as described earlier^90^. Briefly, samples were fixed in 4% PFA at 4 °C overnight. Next, samples were incubated in PBSG-T (0.2% gelatin, 0.01% thimerosal, and 0.5% TritonX100 in PBS) for 3 days with rotation (70 rpm) at room temperature to block non-specific antibody binding. Lung samples then were incubated with an anti-mouse-GFP antibody (clone: ab13970, APCAM, Waltham, USA) which is diluted in 0.1% Saponin in PBSG-T for 7 days with rotation (70 rpm) at 37℃. Afterward, lung samples were washed 6 times with PBST (0.5% Triton in PBS) for 1 hour each and incubated with goat anti-chicken (AF647) secondary antibody (ab150175, APCAM, Waltham, USA) and DAPI diluted in 0.1% Saponin in PBSG-T for 3 days with rotation (70 rpm) at 37℃. Samples were washed 6 times again in PBST for 1 hour each with rotation at room temperature to finish the last step of staining. Subsequently, clearing was performed after dehydration in a concentration gradient of tetrahydrofuran (THF, Sigma, Burlington, MA, USA, 50% v/v tetrahydrofuran/H2O overnight, 50% THF/H2O for 1 h, 80% THF/H2O for 1 h, 100% THF for 1 h, 100% THF overnight, and 100% THF for 1 h) with continuous gentle shaking. Next samples were incubated in dichloromethane (DCM, Sigma, Burlington, MA, USA) for around 30−40 min and eventually immersed in dibenzyl ether (DBE, Sigma, Burlington, MA, USA) at least 2 h prior to imaging. Imaging was performed in dibenzyl ether with an LSFM (Ultramicroscope II, LaVision Biotec) equipped with an sCMOS camera (Andor Neo, Abingdon, United Kingdom) and a 2× objective lens (Olympus MVPLAPO 2×/0.5 NA) equipped with an Olympus MVX- 10 zoom body, which provided zoom-out and -in ranging from 0.63× up to 6.3×. Light sheet images were generated with different magnification factors and a step size of 5-10 μm according to sample size with 470±30 / 640±30 nm ex/em bandpass filters for QDs. Lung tissue autofluorescence was generally scanned with 520±40 / 585±40 nm ex/em filters to show the microstructure of the lungs. 100 ms exposure time and 95% laser power are typically set with LSFM, where the light sheet has a different xy width and numerical aperture (NA) to match the different sample sizes. During the LSFM image acquisition, the samples were immersed in DBE. Imaris 9.1.0 (Bitplane, Belfast, United Kingdom) was used to perform 2D and 3D rendering and image processing.

### Fluorescence-based analysis of QD-NP dose in lung homogenates

Mice were sacrificed immediately after L-IVM experiments by exsanguination and then the lung perfusion was performed following the above protocol. Subsequently, the whole lungs were removed. Quantification of QDs was based on a previously published protocol^94^. Briefly, each lung was immersed in 1 ml tissue lysing solution (Solvable®, Perkin–Elmer, Waltham, USA) at 50 °C for 24 h until complete tissue dissolution. Subsequently, the QD fluorescence intensity of the samples was measured with a spectrofluorometer (ex/em wavelength: 400 nm/655 nm; Safire 2, Tecan, Zurich, Switzerland) and compared to standard curves, which were generated for each QD type by measuring solubilized blank lung samples with known but different amounts of QDs.

### Multiplex Cytokine/Chemokine Analysis

Cytokines and chemokines in BAL fluid were measured using the multiplex bead array system Bio-Plex Pro Mouse Chemokine Assay Panel 31-Plex (#12009159, Bio-Rad Laboratories GmbH), following the manufacturer’s instructions. Data were acquired using a Luminex200 system with BioPlex Manager 6.1 software. Standard curves were fitted using the logistic-5PL regression type. The data was visualized with a heatmap generated by utilizing the statistical programming environment R (v4.4.4) using the heatmap package (v1.0.12).

### Ex vivo whole lung imaging

To obtain a spatial distribution of fluorescent nanomaterials throughout the lungs, epifluorescence imaging with an IVIS system (Lumina II, Caliper/PerkinElmer, USA) was used for (non-tissue cleared) lungs. In short, the isolated lungs were placed on a platform located in the center of the IVIS chamber and imaged with QDs-specific excitation and emission filters (ex/em: 475 nm/ Cy5.5; exposure time: auto; Binning: Medium; F/Stop: 16 and Lamp level: high). The fluorescence/white light images were acquired with the Living Imaging 4.0 software (Caliper, Newton, Massachusetts, USA) to determine fluorescence intensity. Due to the lack of tissue clearing, the obtained fluorescence profile is only an estimate of the QD distribution in the detector-facing, peripheral layer of the lung. Yet, for the spatially uniform aerosol inhalation, this is a good estimate of QD distribution in the lung.

### Transmission Electron Microscopy

Transmission electron microscopic images of lung tissue were acquired as previously described^95^. Briefly, lungs were perfusion-fixed with 2% formaldehyde, 2% glutaraldehyde in 0.1 M cacodylate buffer (pH 7.4) via the left ventricle. Finally, also the lung was filled with fixative via the trachea. After dissection, the samples were fixed for 3 h at room temperature. The tissue was cut into smaller cubes (1 mm^3^) and postfixed in 1% OsO4 containing 1.5% potassium cyanoferrate. After dehydration, the samples were embedded in epon. Ultrathin (70 nm) sections were cut in serial sections (Leica-UC6 ultramicrotome, Vienna, Austria). Except for some sections that were counterstained with uranyl acetate and lead, sections were not stained so that QDs were unambiguously identified by their morphology. The images were acquired at 80 kV on an FEI-Tecnai 12 transmission electron microscope (Thermofisher). Representative areas were imaged with a CCD camera (Veleta, EMSIS, Muenster, Germany).

### MH-S cell culture

Murine MH-S alveolar macrophage-like cells were purchased from the American Type Culture Collection (ATCC®, CRL-2019TM) and grown in RPMI 1640 medium with supplements containing 0.05 mM 2-mercaptoethanol, 10% FBS and 1% penicillin-streptomycin at 37 °C and 5% CO2.

### QD delivery to MH-S cells

500,000 MH-S cells were distributed in 100 μL cell culture medium and then seeded on the apical side of a 6-well transwell insert (4.2 cm^2^, 0.4 μm pores, Corning, New York, USA) placed in a well with 1.7 mL basal medium. At 24 h after cell seeding the basal medium was changed with the aspiration of the 100 µL apical medium, and then the cells were equilibrated at air-liquid interface (ALI) conditions for 2 h. Then the cells were exposed to the QDs aerosol droplets under the ALI conditions using the VITROCELL® Cloud Alpha 6 aerosol-cell exposure system. For this, 100 μl of 1:20 diluted QD suspension in distilled water with 0.1% NaCl (0.7 µg QD/µl) was nebulized with an Aeroneb Pro nebulizer (Aerogen Inc., Ireland) and ca. 5 µl / insert is deposited directly onto the cells^48^. Thereafter, cells were incubated for 2 h and washed with PBS to remove the unattached particles after incubation. IF staining imaging was used for the qualitative comparison of nanoparticle phagocytosis by M-HS across different groups, while FACS was employed for the quantitative analysis of nanoparticle phagocytosis by M-HS in these groups. For IF staining, M-HS cells were fixed with 4% PFA for 10 min, followed by staining with DAPI (1 μg/ml, Roche, Basel, Switzerland) and AF488-labelled phalloidin (5 U/L, A12379, Thermo Fisher Scientific, Carlsbad, USA) for 30 min. The cells were then washed three times with DPBS containing 0.1% Tween 20, each wash lasting 5 min. Finally, the stained samples were mounted using DAKO fluorescence mounting medium (Dako Omnis, Agilent, Santa Clara, USA) and covered with coverslips. For FACS, after washing in the FACS buffer (2% BSA) and 1 mM EDTA in PBS), cells were collected and analyzed using a BD FACSCanto II flow cytometer (BD Bioscience, New Jersey, U.S.) with BD FACSDiva software.

### Water-soluble tetrazolium salt (WST) cell viability assay

MH-S cells were seeded in 24-well plates at a density of 10^5^ / well and cultured overnight. Cells were exposed to QD particles (8nM) and vehicle as a control in 500μl cell culture medium in submerged conditions for 1 h. The supernatant was removed, and the cells were incubated with 10 % WST reagent (Roche, Basel, Switzerland) for 15 min at 37 °C and 5% CO2. Next, WST samples were collected and centrifuged at 14,000 rpm, 10 min at room temperature. Absorbance at 450 nm of 200 µl solution was measured in 96-well plates with a plate reader (TECAN Trading AG, Männedorf, Switzerland). Each sample was measured from two independent wells. The absorbance value was corrected with a blank sample (WST solution non-incubated with cells). The cell viability of each sample was compared with the control group according to the following equation: Cell viability= (sample OD – blank OD)/(control OD – blank OD) *100%.

### Phagocytosis assay

MH-S alveolar macrophage-like cells were seeded in 24-well plates at a density of 25 * 10^4^ cells / well. Cells were pre-treated with anti-ICAM-1, anti-LFA-1, anti-CD44, anti-CD64, anti- CD88 and isotype (10 μg/ml) antibodies, cromolyn (4 μg/ml) or vehicle-control, respectively for 3 h at 37 °C and 5 % CO2 (Table 3). Thereafter pHrodo™ Green E. coli BioParticles (5 μg/ml, Thermo Fisher Scientific, Carlsbad, USA), pH-dependent fluorescent particles, were added to the supernatant evenly for 1h. After washing in the FACS buffer, cells were collected and analyzed using a BD FACSCanto II flow cytometer (BD Bioscience, New Jersey, U.S.) with BD FACSDiva software.

**Table 3:** List of blocking antibodies or inhibitors applied in vitro.

| Blocking antibody/Inhibitor | Clone | Manufacturer |
| --- | --- | --- |
| **In vivo mAb anti-mouse LFA-1α** | M17/4 | Bio X Cell,<br>Lebanon, USA |
| **Purified anti-mouse CD54 (ICAM-1) antibody** | YN1/1.7.4 | ATCC,<br>Manassas, USA |
| **Ultra-LEAF™ Purified anti-mouse CD88 (C5aR) mAb** | 20/70 | BioLegend,<br>Fell, Germany |
| **Ultra-LEAF™ Purified anti-mouse CD64 (FcγRI) mAb** | W18349F | BioLegend,<br>Fell, Germany |
| **In Vivo mAb rat IgG2a isotype control** | 2A3 | Bio X Cell,<br>Lebanon, USA |
| **Ultra-LEAF™ Purified Rat IgG2b, κ Isotype Ctrl antibody** | RTK4530 | BioLegend,<br>Fell, Germany |

### Statistical analysis

All data were presented as Mean ± standard error of the Mean (SEM) and plotted with GraphPad Prism 8 (GraphPad Software Inc., La Jolla, USA), with the sample sizes and the number of repeats indicated in the figure legends. Comparison of results between two groups for normally distributed data were analyzed using the two-sided student t-test and for nonparametric data with the Mann-Whitney rank-sum test. Comparisons among multiple groups were performed using a one-way ANOVA with Tukey’s comparisons test.

Significances are defined as 0.05 (p < 0.05, *), 0.01 (p < 0.01, **), 0.001 (p < 0.001, ***) and 0.0001 (p < 0.001, ****), while p-value ≥0.05 was considered not significant (ns).

## Acknowledgements

The authors would like to particularly thank Dr. Annette Feuchtinger (Research Unit Analytical Pathology, Helmholtz Munich) for support in light sheet imaging.

## Author contributions

QL and MR designed and planned the project. QL performed all the experiments and analyzed the data. LY provided support on animal experiments and image analysis. CL and QZ provided support on in-vitro experiments. LH provided support on figure revision. AS and OS designed and integrated the nebulizer inhalation system. DK provided support on animal experiments. DZ performed Transmission Electron Microscopy. JS, AÖI and LC provided critical tools. QL, TS and MR wrote the manuscript. QL, LY, LC, OS, MS, TS and MR revised the manuscript. All authors read, discussed, improved, and approved the publication. Correspondence to QL and MR.

