## Supplemental Information for "Alveolar macrophages initiate the spatially targeted neutrophil recruitment during nanoparticle inhalation"

### Supplemental figures

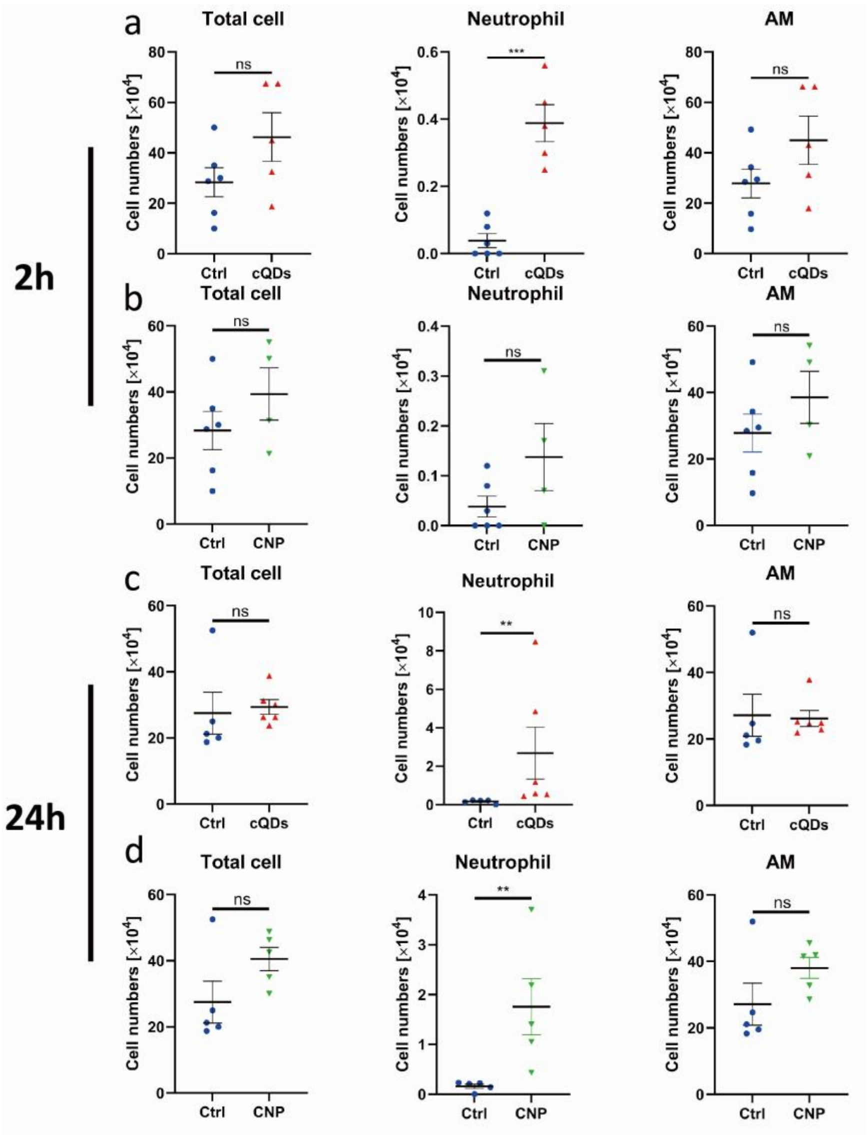


**Figure S1. Bronchiolar-alveolar-lavage (BAL) cell numbers at different time points (2 h and 24 h) after cQDs/CNP inhalation**

(a) and (b) 2 h and (c) and (d) 24 h after NP inhalation the numbers of total cells, AMs and neutrophils were determined in May Grunwald-stained-cytospin samples of cQD and CNP exposed mice (16 $cm^2/g$ cQDs; 170 $cm^2/g$ CNP), respectively, and compared to vehicle controls. Data are presented as means ± SEM, n= 4-6 mice/group, ns: $p \geq 0.05$, *: $p < 0.05$, **, $p < 0.01$ and ***, $p < 0.001$ by paired Student's t-test (parametric data) or Mann-Whitney rank-sum test (nonparametric data).

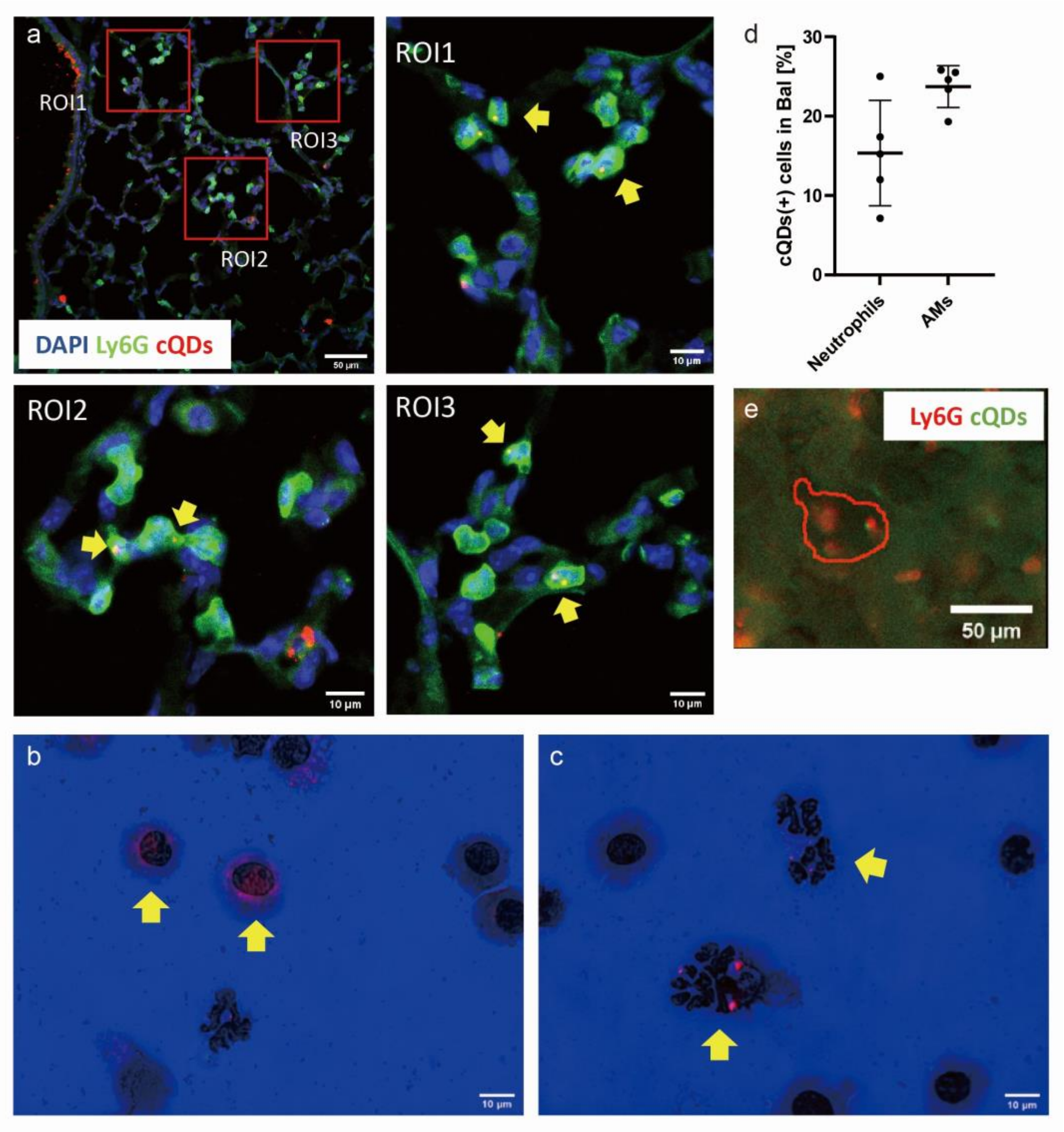


**Figure S2. Infiltrating neutrophils internalize nanoparticles in the alveoli**

(a) Fluorescence confocal microscopy of WT mice lung slices obtained 90 min after cQDs (red) inhalation. Neutrophils labeled with anti-Ly6G mAb (green). Nuclei stained by DAPI (blue) (Scale bar: 50 µm). Frequently alveolar localized neutrophils are colocalized with cQD fluorescent spots, indicating cQDs cellular uptake. ROI1-ROI3: Magnifications highlight cQDs-laden neutrophils (arrows), scale bars: 10 µm.

(b) and (c) cQDs (red) can be detected in AMs (b, arrows) as well as in neutrophils (c, arrows) in May Grunwald-stained BAL cytospin-samples, collected at 24 h after cQD exposure, by combined epifluorescence and phase contrast imaging (Scale bars: 10 µm).

(d) Quantification of cQD positive AMs and neutrophils in BAL samples obtained 24 h after cQDs inhalation, n=4 mice.

(e) L-IVM image at 60 min after cQDs-inhalation shows cQD colocalization with alveolar localized neutrophils (cQDs: green, neutrophils labeled by anti-Ly6G mAbs (i.v): red, scale bar: 50 µm).

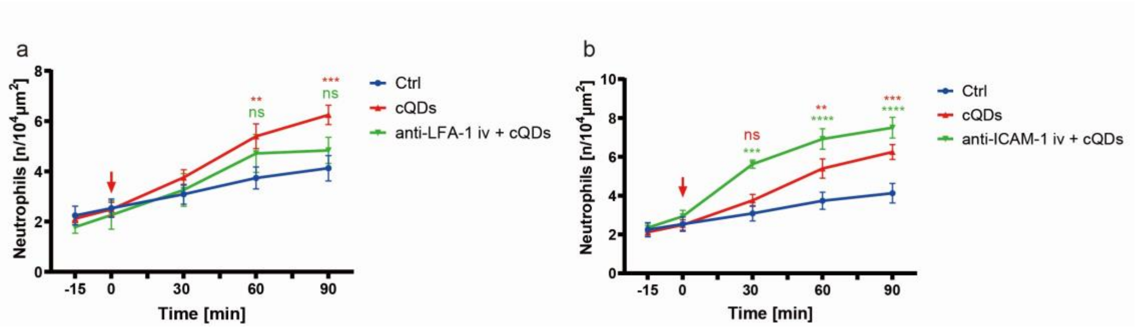


**Figure S3. Blockade of intravascular LFA-1 as well as ICAM-1 has no mitigating effect on cQDs-evoked recruitment of neutrophils**

(a)-(b) Alterations in neutrophil numbers in the alveolar region over 90 min were analyzed by L-IVM in mice receiving LFA-1 or ICAM-1 blocking mAbs via intravenous injection 30 min prior to inhalation of cQDs or vehicle-control, n=3 mice/group. Data are presented as Means ± SEM. ns: $p \geq 0.05$, *: $p < 0.05$, **: $p < 0.01$, ***: $p < 0.001$ and ****: $p < 0.0001$.

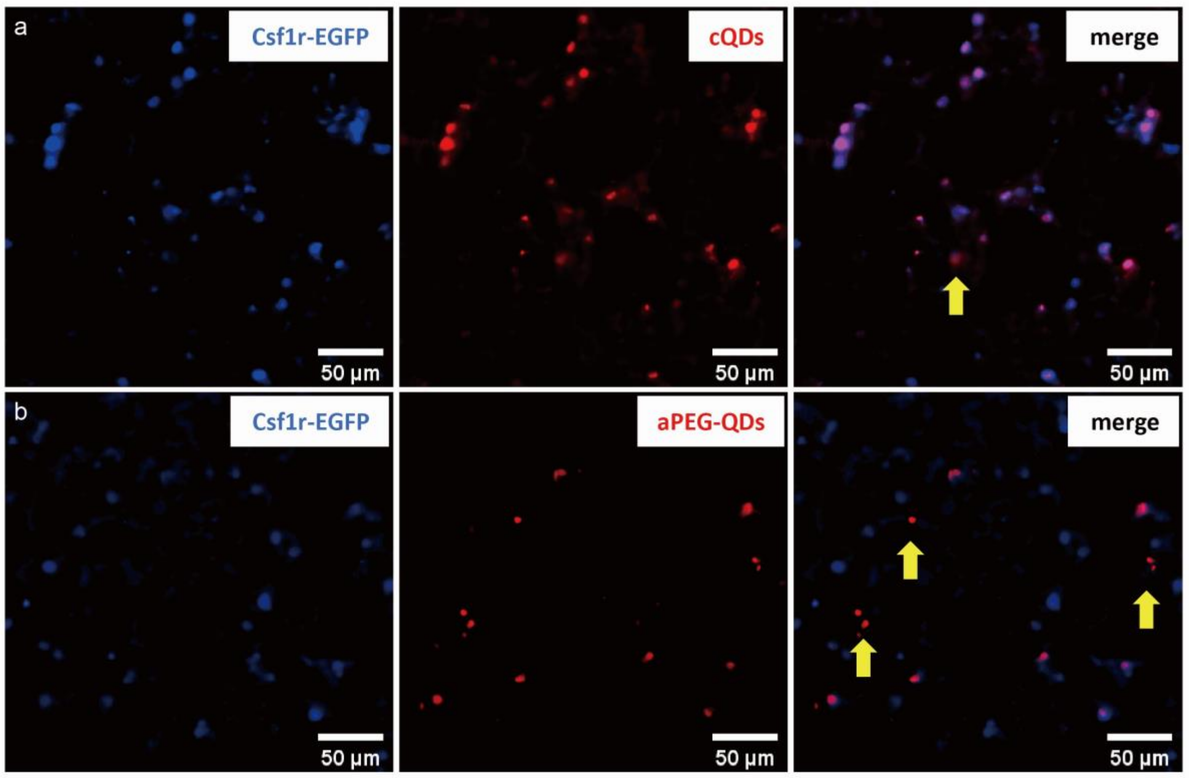


**Figure S4. PEGylation prevents QD NPs from being internalized by AMs**

(a)-(b) Tissue clearing was used to achieve co-imaging of AMs from Macgreen mice (Csf1r-EGFP is expressed selectively in macrophage and monocyte cell lineages) after transcranial perfusion for flushing out all Csf1r-EGFP labeled cells in the blood. 2D images from lightsheet fluorescence microscope images of cQDs and aPEG-QDs exposed mice (2 h). Csf1r-1(+) AMs are shown in blue, QDs are shown in red, AMs with ingested QDs ingested appear in purple. (panel a: cQDs, panel b: aPEG-QDs, Scale bar: 50 µm).

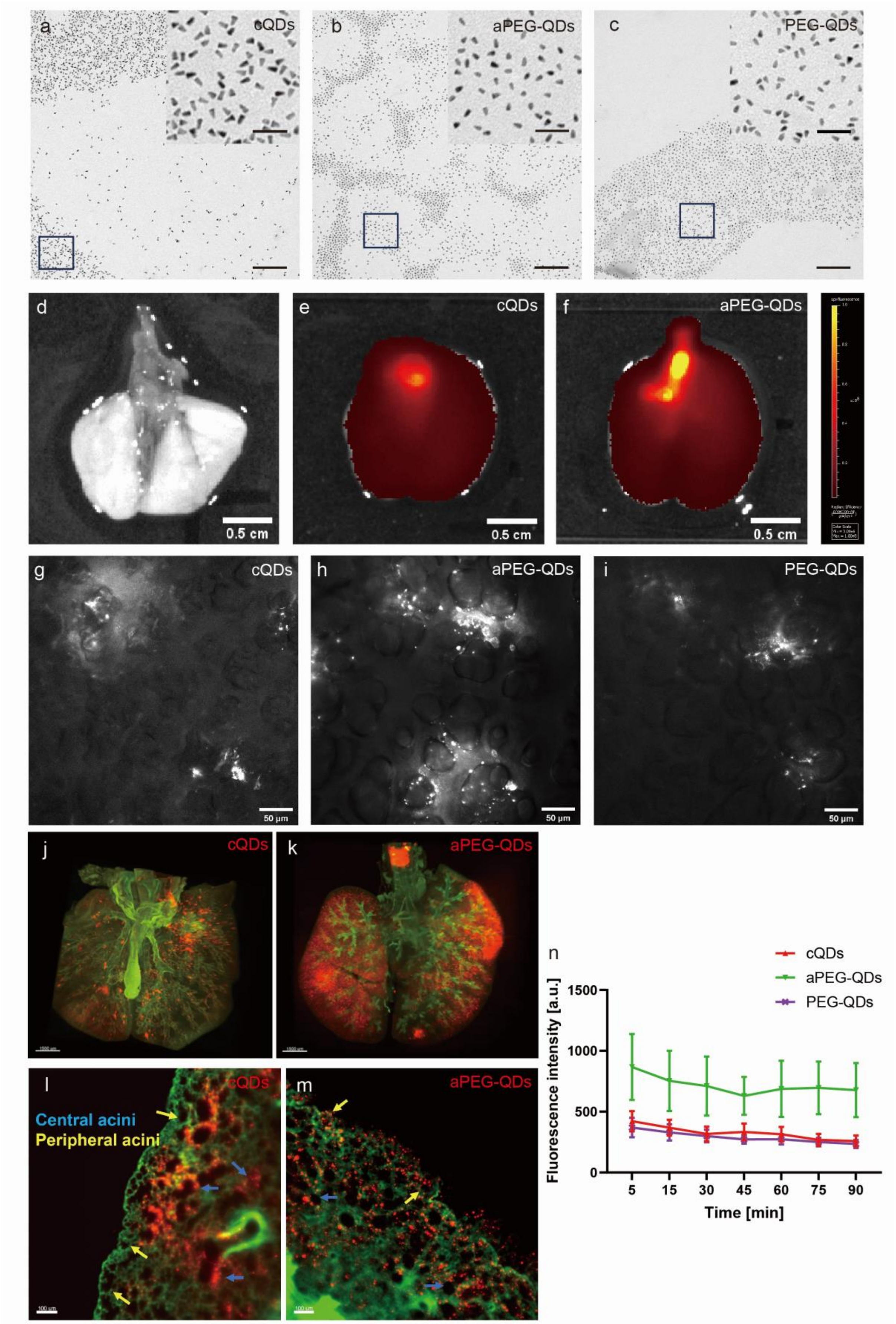


**Figure S5. 3D/2D mapping of QD distribution in entire mouse lungs upon ventilator-assisted QD-NP aerosol inhalation**

(a)-(c) Electron microscopic images of the three different QD-species. QDs were diluted in distilled water (panel a: cQDs, panel b: aPEG-QDs and panel c: PEG-QDs, scale bars: 200 nm).
(d)-(f) Analysis of cQD and aPEG-QD distribution in whole lungs using epifluorescence imaging. Typical ex vivo lung images from mice at 2 h after receiving 16 $cm^2/g$ (geom-surface area of NPs / mass-lung) cQDs or aPEG-QDs via inhalation and vehicle control using the IVIS system, which is particularly sensitive to the most peripheral part of the lung which is the focus of L-IVM analysis (panel d: Ctrl, panel e: cQDs, panel f: aPEG-QDs).
(g)-(i) L-IVM images of the three types of QDs (g: cQDs; h: aPEG-QDs; i: PEG-QDs) showing the QD distribution pattern in the most peripheral alveolar part of the lungs 5 min after pulmonary NP inhalation. QDs are shown in white. Scale bars: 50 µm.
(j)-(m) The distribution of QDs (red) in the Z-stack images of entire cleared lungs obtained by light sheet microscopy (panel j and k: 3D MIP, l: cQDs and m: aPEG-QDs, scale bar: 1500 µm) and in single z-planes (panel l and m: 2D xy slice, l: cQDs and m: aPEG-QDs, scale bar: 200 µm) of the tissue structure (autofluorescence, green) 1h after inhalation of different types of (16 $cm^2/g$) QDs.
(n) Average QD fluorescence intensities in L-IVM images at consecutive time points after inhalation of cQDs, aPEG-QDs, or PEG-QDs (Means ± SEM, n= 3 mice/group)

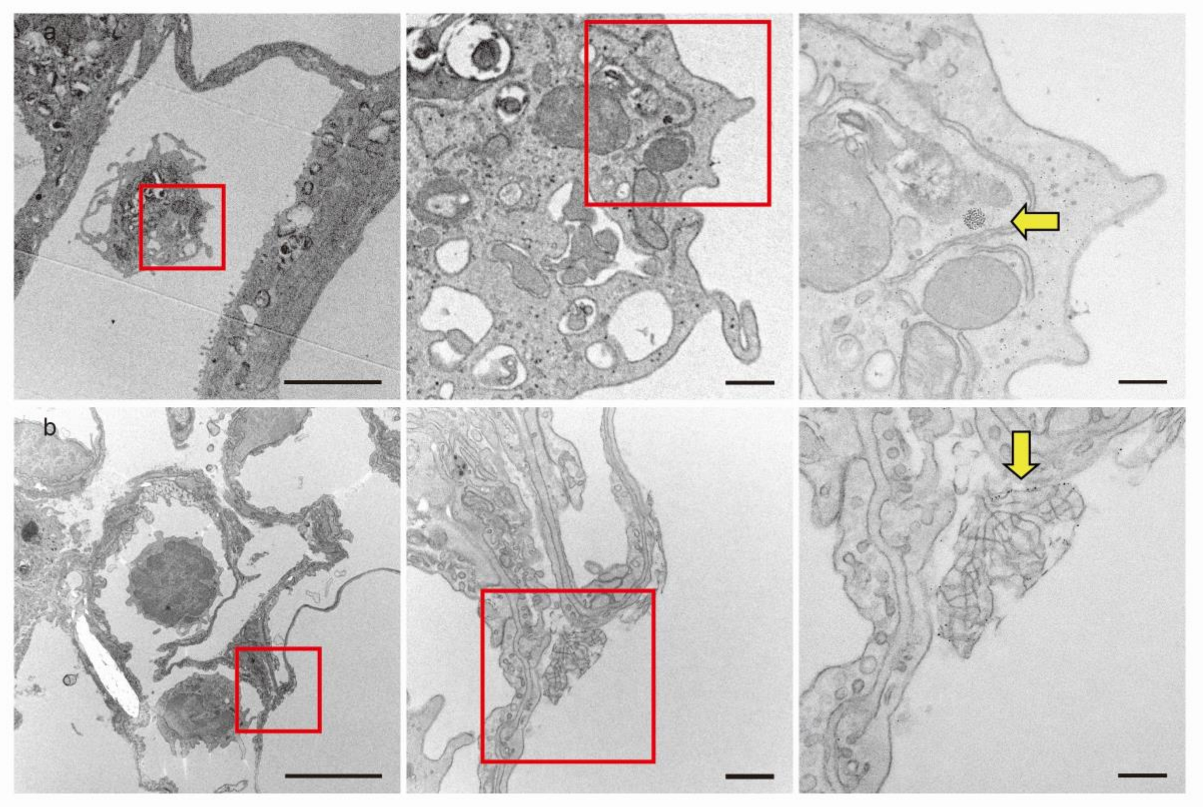


**Figure S6. cQDs accumulate in AMs and are attached to surfactant lattices**

Transmission electron microscopy of cQDs-exposed lungs (1 h). (a) Aggregates of cQDs were found within the cytoplasm and endolysosomes of AMs. (b) Non-cell associated “free” cQDs can also be found attached to surfactant components (tubular myelin sheaths) at the alveolar air-liquid interface. QDs are visible as distinct black points (yellow arrows) within the surfactant lattice. Overview and detailed higher magnification pictures of depicted areas with corresponding scale bars: 5 µm, 500 nm, 200 nm.
